# Molecular characterization of Left-Right symmetry breaking in the mouse embryo

**DOI:** 10.1101/2022.12.26.521947

**Authors:** Richard C.V. Tyser, Ximena Ibarra-Soria, Monique Pedroza, Antonio M.A. Miranda, Teun A.H. van den Brand, Antonio Scialdone, John C. Marioni, Shankar Srinivas

## Abstract

The asymmetric morphology of the mammalian heart is essential to its function as the organ of pulmonary and systemic double circulation. Left-right asymmetry is established by a leftward flow in the node that results in the asymmetric expression of *Nodal*. This triggers a cascade of asymmetric expression of downstream genes, such as *Pitx2c*, in the lateral plate mesoderm that gives rise to the first morphologically recognizable primordial heart structure, the cardiac crescent. Relatively little is known about gene expression asymmetries in the cardiac crescent that might underpin asymmetric cardiac morphogenesis. To systematically identify asymmetrically expressed genes, we performed a single-cell transcriptional analysis of manually dissected left and right halves of the cardiac crescent at stages spanning symmetry breaking. This revealed both left and right-sided genes that have not previously been implicated in left-right symmetry breaking. Some of these were expressed in multiple cell types but showed asymmetric expression in only a sub-set of cell types. We validated these findings using multiplexed *in situ* Hybridization Chain Reaction (HCR) and high-resolution volume imaging to characterize the expression patterns of select genes. Using *Dnah*^*iv/iv*^ mutant embryos that show randomized situs, we established that all the genes tested tracked the asymmetric expression of *Pitx2c*, indicating their asymmetric expression also arose from the early asymmetries at the node. This study provides a high-fidelity molecular characterization of left-right symmetry breaking during cardiac crescent formation, providing a basis for future mechanistic studies on asymmetric cardiac morphogenesis.

## INTRODUCTION

The vertebrate body plan is asymmetric, with most internal organs being asymmetrically shaped and positioned along the left-right (LR) body axis. These differences arise due to the regulation of several conserved genes early in development, around embryonic day (E) 7.5. Early bilateral symmetry breaking is initiated by the node1. The node is comprised of ciliated cells, which generate a leftward fluid flow that leads to the asymmetric activation of *Nodal*, a TGFβ growth factor, in the lateral plate mesoderm (LPM)^2–5^. Transient activation of *Nodal* results in the induction of transcription factors *Lefty* and *Pitx2*, which play a fundamental role in governing asymmetric organ morphogenesis^6–11^. Of the different *Pitx2* isoforms, it is the *Pitx2c* isoform that is asymmetrically expressed and, in contrast to transient *Nodal* expression, is maintained asymmetrically throughout organogenesis^12,13^. *Pitx2c* is the main cardiac isoform and is dynamically expressed during heart formation^14^.

Heart formation is an excellent model to investigate LR differences as it is the first organ to form and function as well as display morphological asymmetries around E8.25^15^. In humans, defects in left-right asymmetry patterning lead to a range of congenital heart defects^16^. In the mouse, the heart initially forms as a symmetrical arc of cells, termed the cardiac crescent (CC), at around 7.75dpc before transitioning into the linear heart tube (LHT) which undergoes rightward looping prior to the formation of the four-chambered heart. The process of rightward looping and generation of a helical shape are modulated by *Nodal* at the poles of the LHT, which regulate proliferation, differentiation, and extracellular matrix genes^17^. Detailed morphological analysis and modeling of CC to LHT development has revealed that LR symmetry breaking can already be observed prior to rightward looping of the LHT, with LR differences detected in the orientation of the inflow tracts^18^.

Whilst the morphological differences during heart looping have been well characterized, the early molecular basis is still relatively unclear^17,19,20^. We therefore sought to explore the temporal dynamics of LR molecular differences at the single-cell whole transcriptome level during cardiac crescent development and the onset of embryo wide symmetry breaking. To do this we exploited a single-cell transcriptomic dataset of the developing cardiac crescent that we had previously generated, in which we collected cells separately from the left and right cardiac regions. This dataset covers the period of development in which LR symmetry breaking is first established and allows the identification of novel asymmetrically expressed genes, thereby expanding our molecular understanding of how asymmetries arise in the forming embryo.

## RESULTS

### Identifying genes expressed asymmetrically in the forming cardiac crescent

Previously, we generated an unbiased single-cell transcriptomic dataset of the developing mouse heart, from emergence of the cardiac crescent to the formation of the linear heart tube, accessible at https://marionilab.cruk.cam.ac.uk/heartAtlas/ ^21^. We identified twelve different cell populations, from all three germ layers. These included mesodermal cell types (cardiac progenitors, cardiomyocytes, endothelial and blood populations), overlying endoderm (including definitive and yolk sac) and adjacent ectoderm (representing surface/amnion and neuroectoderm)^21^. To discover left-right asymmetrically expressed genes, we manually dissected cells from the left or right sides of the embryos during the collection of cells from the cardiac crescent (Figure 1a). Thus, these data provide a highly time-resolved resource to investigate asymmetrically expressed genes during the onset of left-right asymmetry in the heart. Across the four stages of cardiac crescent development profiled, we generated high-quality transcriptomes for roughly equal numbers of cells from the left and right sides (Figure 1b). Most of the twelve cell subpopulations also had equal representation of cells from both sides, and these intermingled in the UMAP representation (Figure 1c, S1a). To confirm our dissection and faithful separation of the left and right cardiac crescent regions, we first examined the well-known asymmetrically expressed genes *Nodal, Lefty* and *Pitx2* (Figure 1d and S1b)^2,3,6–11^. These three genes were predominantly expressed in the cardiac mesoderm related clusters (Me3-Me8), representing both cardiac progenitors and cardiomyocytes, and as expected, were expressed at significantly higher levels and in more cells from the left (Figure 1d and S1b). Cell cycle analysis showed no difference in the proportion of cells in G2/M, G1 or S-phase between left and right sides of the embryo in cardiomyocytes (Me3), blood (Me1), endothelial cells (Me2), endoderm (En1, En2) and 2) or ectoderm (Ec1, Ec2) although we observed a small but significant reduction of cells in G1 phase from the right side in the cardiac mesoderm progenitor clusters (Me4-7) (Figure 1e, S1c, Supplementary Table 1).

**Figure 1:**
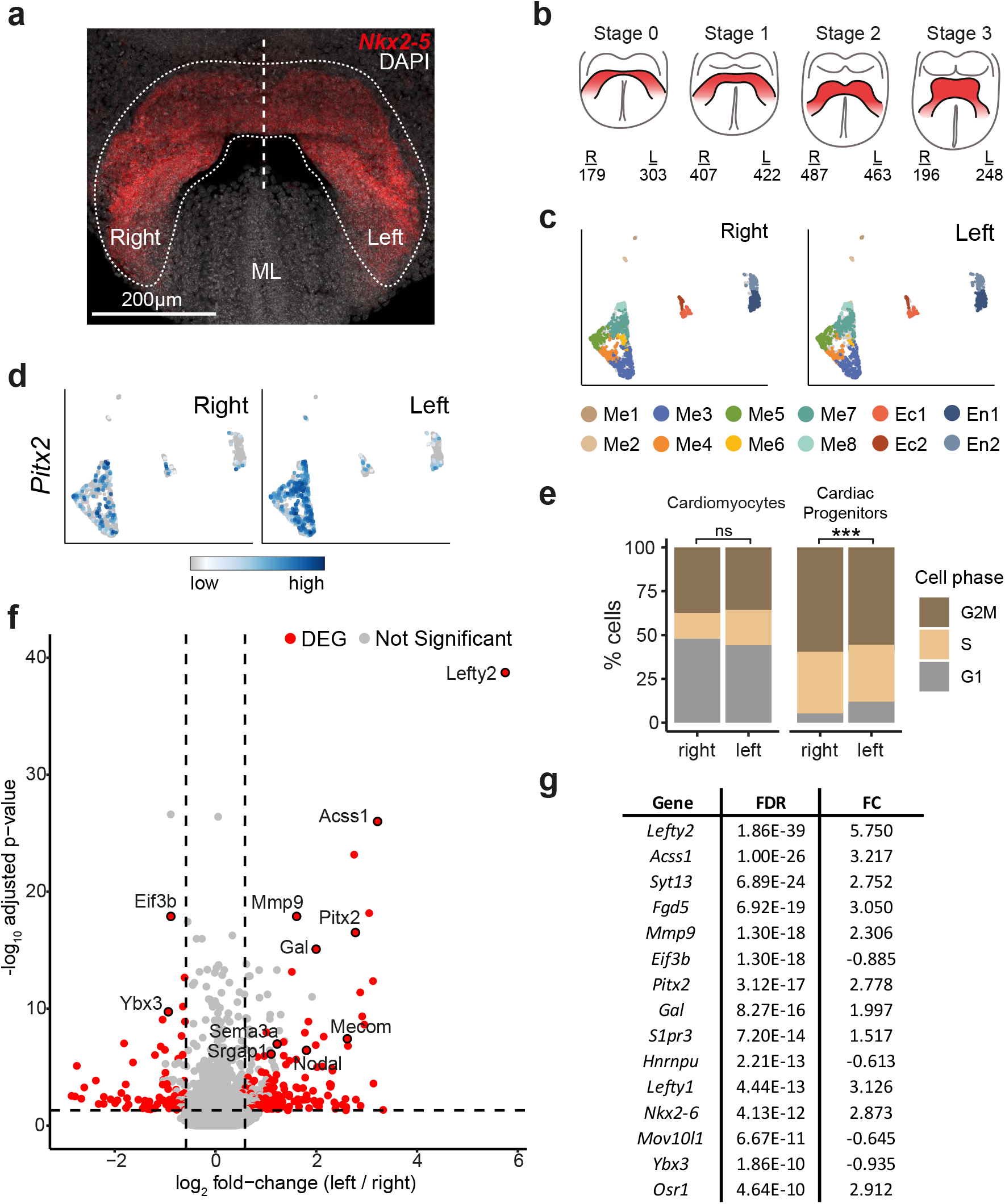
Single-cell molecular characterization of left right differences during cardiac crescent development. **a**, Ventral maximum intensity projection (MIP) of a stage 1 cardiac crescent stage embryo. The cardiac crescent is marked by NKX2-5 protein expression. The dotted region highlights the area micro-dissected for collecting cells and the L/R separation. ML, midline. **b**, Schematic of crescent stages collected for single-cell RNA-seq highlighting the number of cells collected from either the left or right that passed quality control. **c**, UMAP plot of all cells collected from either the left or right that passed quality control (n = 2,705) computed using highly variable genes. Cells with similar transcriptional profiles were clustered into 12 different groups, as indicated by the different colors, and were equally represented in both the left and the right. **d**, UMAP showing the expression of the well-characterized left-sided marker gene *Pitx2* in cells collected from either the left or right side of the forming cardiac crescent. **e**, Bar graphs showing the proportion of left and right cells from cardiomyocytes (Me3) and cardiac progenitors (Me4-Me7) in either G1, S or G2M phase of the cell cycle (binomial test that the proportion of cells in G1 is lower in the right side; ***p<0.001; ns = not significant). **f**, Differential expression analysis in mesodermal cell clusters (Me3-Me8). The adjusted p-value (y-axis) is plotted against the fold-change between left-right cells (x-axis). For non-differentially expressed genes (grey), the average fold-change for all clusters is used; for DEGs (red) the largest fold-change (in clusters where the gene is expressed) is used. Genes are considered DE if their adjusted p-value is lower than 0.05 and their fold-change is larger than 1.5 (indicated by the dashed lines) in at least one cluster. **g**, Table showing the top 15 DEGs, along with false discovery rate (FDR) and largest (absolute) log2 fold change (FC) when comparing left to right sided cells.

To identify novel asymmetrically expressed genes, we used differential expression analysis to test for expression differences between the left and right cells in each cell subpopulation (excluding Me1 and Me2 due to their small number of cells). In the mesodermal cell types, we identified 229 asymmetrically expressed genes, with two thirds expressed at higher levels in the cells from the left side of the embryo (Figure 1f and g, Supplementary Table 2). As observed before, *Lefty2* and *Pitx2* were among the most significant asymmetrically expressed genes, as well as *Atp1a1* and *Gal*, which has also been reported to be expressed at higher levels in the left side of the embryo^22^. The endodermal and ectodermal subpopulations showed fewer significant differences (23 and 60 respectively; Figure S2a and b, Supplementary Table 2). There was almost no overlap between the differentially expressed genes identified in each germ layer (only six genes were differentially expressed in the mesoderm clusters that were also differentially expressed in the endoderm (*Eif3b, Polr2a, Gm13050*) or in the ectoderm (*Gm18821, Efhd2, Klf6*), and none in all three.

The genes with asymmetric left-right expression in cardiac mesoderm cells (Me3-Me8) were expressed in varying patterns across the different clusters. Most DE genes were preferentially expressed either in the most mature cardiomyocytes (Me3; 58/229 genes) or in SHF progenitor cells (Me6-Me8; 72/229 genes; Supplementary Table 2). Interestingly, the genes with expression in the SHF-related clusters showed significantly larger fold-changes and smaller p-values compared to the genes in other clusters.

### Validation of asymmetric expression using HCR

To experimentally validate the asymmetric expression of these genes, we used multiplexed Hybridization Chain Reaction (HCR) and confocal volume imaging. We used *Nkx2-5* and *Pitx2c* expression as positive controls for symmetrical and asymmetrical cardiac progenitor expression respectively. Multiplexed detection of these controls along with the transcript being tested allowed us to circumvent confounders that may have resulted from dynamic changes in expression patterns of these genes during development within even closely staged multiple samples.

To effectively use *Pitx2c* as a control for asymmetric expression, we first performed a detailed time-resolved characterization of its expression during early cardiac development stage mouse embryos. *Pitx2c* is the left specific isoform of *Pitx2* and has previously been reported to be expressed in the heart, gut and LPM^12,13^. To establish when and where asymmetric expression of *Pitx2c* first occurs during cardiac crescent formation, we used whole mount multiplexed in situ HCR, followed by confocal volume imaging (Figure 2a). This showed that in the majority of late headfold stage embryos^23^ (stage -1 cardiac crescents) *Pitx2c* was LR symmetrically expressed (Figure 2a) in all tissues (10/12 embryos). *Pitx2c* transcripts were detected in the cranial mesoderm, yolk sac endoderm, paraxial mesoderm and headfold ectoderm (Figure 1d, 2a and S3a-c). In a minority of embryos (2/12), *Pitx2c* was detected asymmetrically in a small region of left caudal LPM, closest to the node and furthest from the cardiac crescent. At stage 0, *Pitx2c* expression remained symmetrical in the paraxial and cranial mesoderm, headfold ectoderm and yolk sac endoderm (Figure 2a). However, by this stage, *Pitx2c* was detected asymmetrically in left-sided LPM in a majority of embryos (7/8), either in a small left-sided region of caudal LPM (4/7 embryos) or had begun to spread rostrally throughout the left-sided LPM extending into the maturing cardiac crescent (3/7 embryos) (Figure 2a and S3d-i). By stage 1, most embryos (29/30) showed left-asymmetric expression throughout the LPM. Symmetric *Pitx2c* expression was still maintained in the headfold ectoderm, cranial mesoderm and yolk sac endoderm, but was lost from the paraxial mesoderm below the crescent (Figure 2a and S4). Once the linear heart tube had formed, asymmetric *Pitx2c* expression was restricted to the left sinus horn and LPM, whilst symmetric expression was maintained in the headfolds, cranial mesoderm and yolk sac. This reveals that the expression of *Pitx2c* is extremely dynamic over the approximately 10-hour period spanning stages -1 to LHT (Figure 2b).

**Figure 2:**
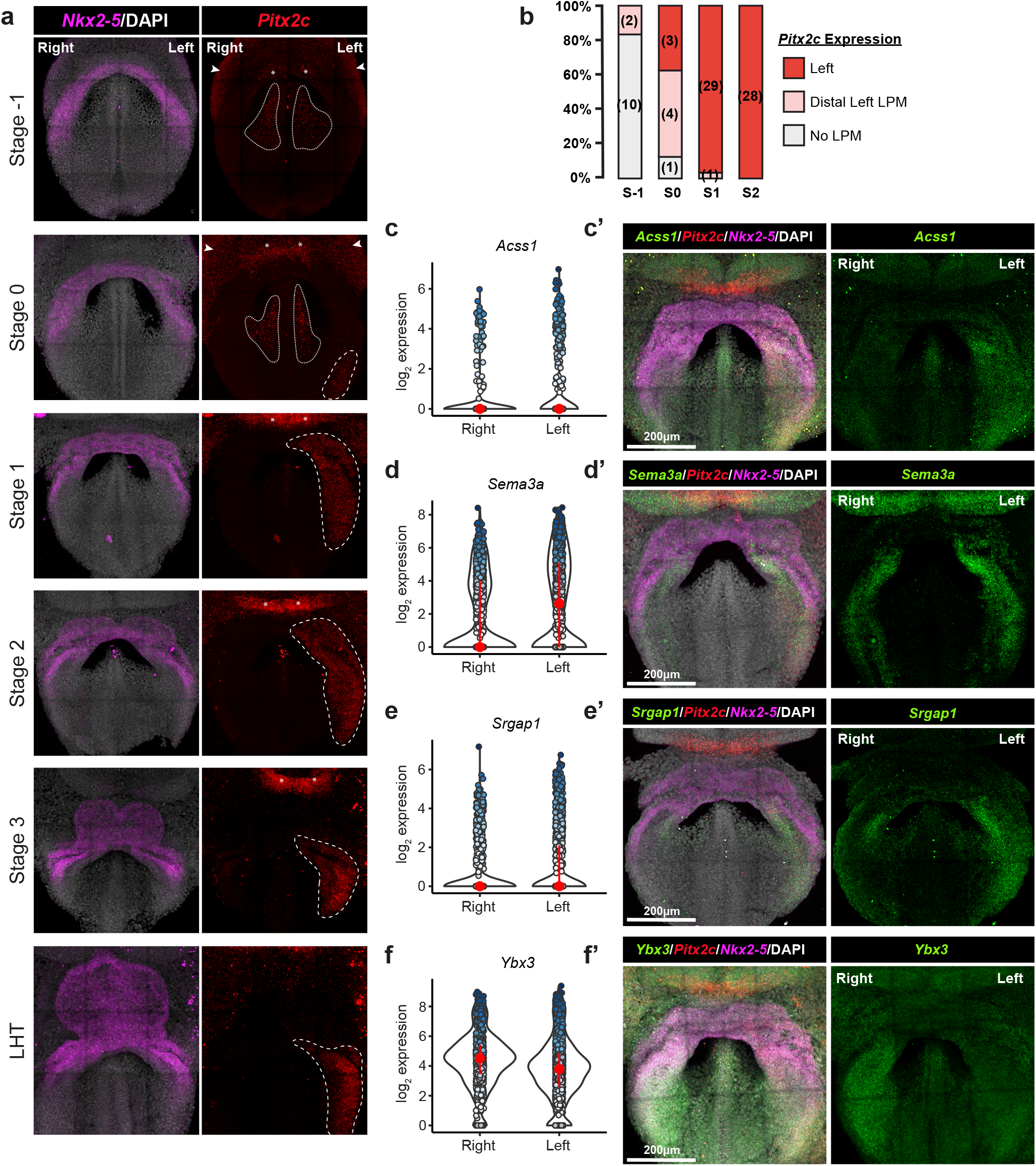
Identification and validation of asymmetrically expressed genes. **a**, MIPs covering stages of cardiac crescent development showing expression of *Nkx2-5* and *Pitx2c* using Hybridization Chain Reaction. *Pitx2c* expression is detected in the lateral plate mesoderm (dashed regions), paraxial mesoderm (dotted regions), yolk sac endoderm (arrowheads) and ectoderm (asterisks) at varying stages of crescent development. **b**, Quantification of asymmetry in *Pitx2c* expression at different stages of cardiac crescent development. A total of 78 embryos were quantified across stages -1 to 2 of crescent development; numbers per stage shown in brackets. **c-f**, Violin plots showing the expression of *Acss1, Sema3a, Srgap1*, and *Ybx3* in left and right mesodermal cells. Darker blue dots represent single cells with higher expression. **c’-f’**, Maximum Intensity Projection (MIP) of hybridization chain reaction (HCR) staining revealing the expression pattern of selected genes as well as *Pitx2c* and *Nkx2-5* as controls, thus validating computational observations. Individual channels are shown in Supplementary Figure 5.

Next, we selected five left-sided genes identified from the single-cell RNA-seq dataset to test: *Acss1, Mmp9, Mecom, Sema3a* and *Srgap1*. HCR analysis was initially performed on mid cardiac crescent stage (stages 1–3) embryos, when *Pitx2c* was clearly asymmetrically expressed. All the left-sided genes tested showed left-sided asymmetry, but their expression profiles differed. *Acss1*, which encodes a mitochondrial acetyl-CoA synthetase enzyme, was predominantly expressed in the left LPM (10/10 stage 1–3 embryos), extending into the cardiac crescent, with diffuse expression throughout this region (Figure 2c, S5 and S6a). In contrast, *Sema3a*, encoding a member of the semaphorin signaling family, and *Srgap1*, encoding the SLIT-ROBO Rho GTPase Activating Protein 1, were expressed in both the left and right LPM extending into the cardiac crescent, but in all cases, more strongly on the left (*Sema3a*; 6/6 stage 1–3 embryos, *Srgap1*; 7/8 stage 1–3 embryos) (Figure 2d, 2e, S5 and S6a). *Sema3a* also showed strong symmetrical expression within the headfolds, consistent with its expression in the transcriptional clusters Ec1 and 2 (Figure S6a).

*Mecom*, which encodes the transcription factor EVI1, was identified in two mesodermal clusters, Me2 (endocardium) and Me6 (differentiating cardiac progenitors). *Mecom* was symmetrically expressed in cluster Me2 and asymmetrically expressed in cluster Me6 (Figure 3a). HCR validated this and showed strong left-asymmetric expression in the LPM (9/9 stage 1–3 embryos) but symmetrical expression in the forming endocardial cells within the cardiac crescent (Figure 3b, 3c and S6a). *Mmp9*, which encodes a matrix metalloprotease involved in extracellular matrix remodeling, was detected within individual cells in the left limb of the cardiac crescent (12/12 stage 1–3 embryos) (Figure 3d, 3e and S6). At slightly later LHT stages, *Mmp9* continues to be asymmetrically expressed and is downregulated in *Nodal* mutants^17^. The anatomical characterization and validation of left-sided expression of *Acss1, Mmp9, Mecom, Sema3a, Srgap1* confirms our single-cell transcriptional approach to identify novel asymmetrically expressed genes.

**Figure 3:**
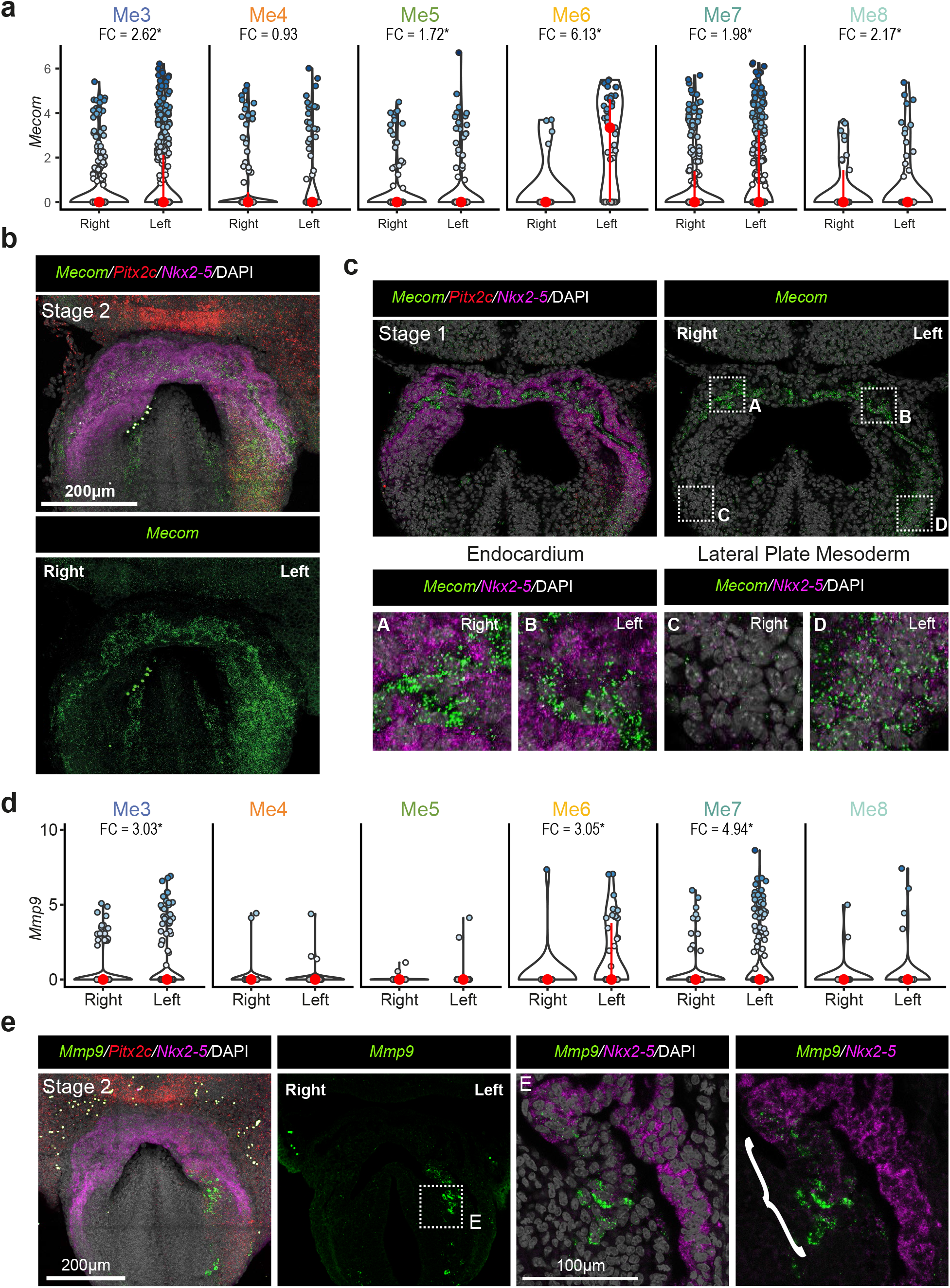
Asymmetrical gene expression varies depending on mesodermal cell type. **a**, Violin plots showing the expression of *Mecom* in the different mesodermal clusters. Darker blue dots represent single cells with higher expression. Changes were considered significant if the fold change (in brackets) was above 1.5 but only for clusters in which the gene was detected at moderate levels. **b**, Maximum Intensity Projection (MIP) of a hybridization chain reaction (HCR) staining revealing the expression pattern of *Mecom, Pitx2c* and *Nkx2-5* in a stage 2 cardiac crescent embryo. **c**, Single Z section of a HCR staining for *Mecom, Pitx2c* and *Nkx2-5* highlighting the symmetrical expression of *Mecom* in the forming endocardium (Me2; insets A and B) and asymmetrical expression in the lateral plate mesoderm (Me3, Me5-8; insets C and D). **d**, Same as a, but for *Mmp9*. **e**, MIP of a HCR staining revealing the expression pattern of *Mmp9, Pitx2c* and *Nkx2-5* in a stage 2 cardiac crescent embryo. Single Z section (inset E) showing the expression of *Mmp9* in single cells corresponding to progenitor clusters Me6 and Me7 (bracket).

Our computational analysis also revealed right-sided genes (Figure 1f and 1g), though fewer than left-asymmetric genes. We sought to experimentally validate *Ybx3* and *Eif3b* by HCR. *Ybx3* encodes a DNA-and RNA-binding protein which can regulate transcription and mRNA stability, having a role in proliferation and amino acid uptake^24^. *Ybx3* was expressed in all germ layers including all mesoderm cell types, however it showed significantly greater expression in mesodermal cells collected from the right side (Figure 2f, S5 and S6a). Whole mount HCR imaging revealed the widespread expression of *Ybx3* in tissues derived from all germ layers. Furthermore in 6/9 embryos, it was expressed more strongly in the right LPM compared to the left, whilst the remaining embryos showed symmetrical (2/9) or left-sided expression (1/9) (Figure 2f). We also assessed the right-sided expression of *Eif3b* that encodes the RNA-binding component of the eukaryotic translation initiation factor 3 complex required for protein synthesis initiation. *Eif3b* was strongly expressed in all cell clusters, and this was supported by HCR analysis which revealed strong expression throughout all tissues (Figure S4). Right-sided *Eif3b* expression was detected in 3 out of 6 embryos, with the remaining embryos displaying left-right symmetrical expression.

### Timing of origin of asymmetries in gene expression

To address the timing with which asymmetrical expression of these genes manifested during the emergence of left-right asymmetry, we performed HCR at earlier stages. We also measured *Pitx2c* expression, to determine whether asymmetric expression of these novel genes preceded or followed that of *Pitx2c*. This revealed that all the markers tested showed asymmetric expression only after *Pitx2c* was upregulated in caudal left-side of the embryo (Figure 4). *Acss1* was not expressed at all prior to asymmetric *Pitx2c* expression. *Mmp9* expression was detected in individual cells located symmetrically on both the left and right sides of the embryo at stages when *Pitx2c* was starting to be expressed asymmetrically in the left LPM. As previously reported, *Mecom* had two different domains of expression. As asymmetric *Pitx2c* expression became established, endothelial *Mecom* expression was detected symmetrically but there was no expression in the LPM. At a stage when *Pitx2c* was starting to be expressed asymmetrically in the left LPM, *Srgap1* was expressed symmetrically in the LPM, more rostrally than the asymmetric LPM *Pitx2c* expression. At this stage, *Sema3a* was strongly expressed in both the left and right sides of the neural ectoderm as well as the LPM running caudally from the cardiac crescent (Figure 4). Whilst there was some overlap in the expression of *Sema3a* and *Nkx2-5* at the boundary of the cardiac crescent, *Sema3a* was expressed only at low levels in the cardiac crescent as well as in the cranial and paraxial mesoderm (Figure 4 and S7).

**Figure 4:**
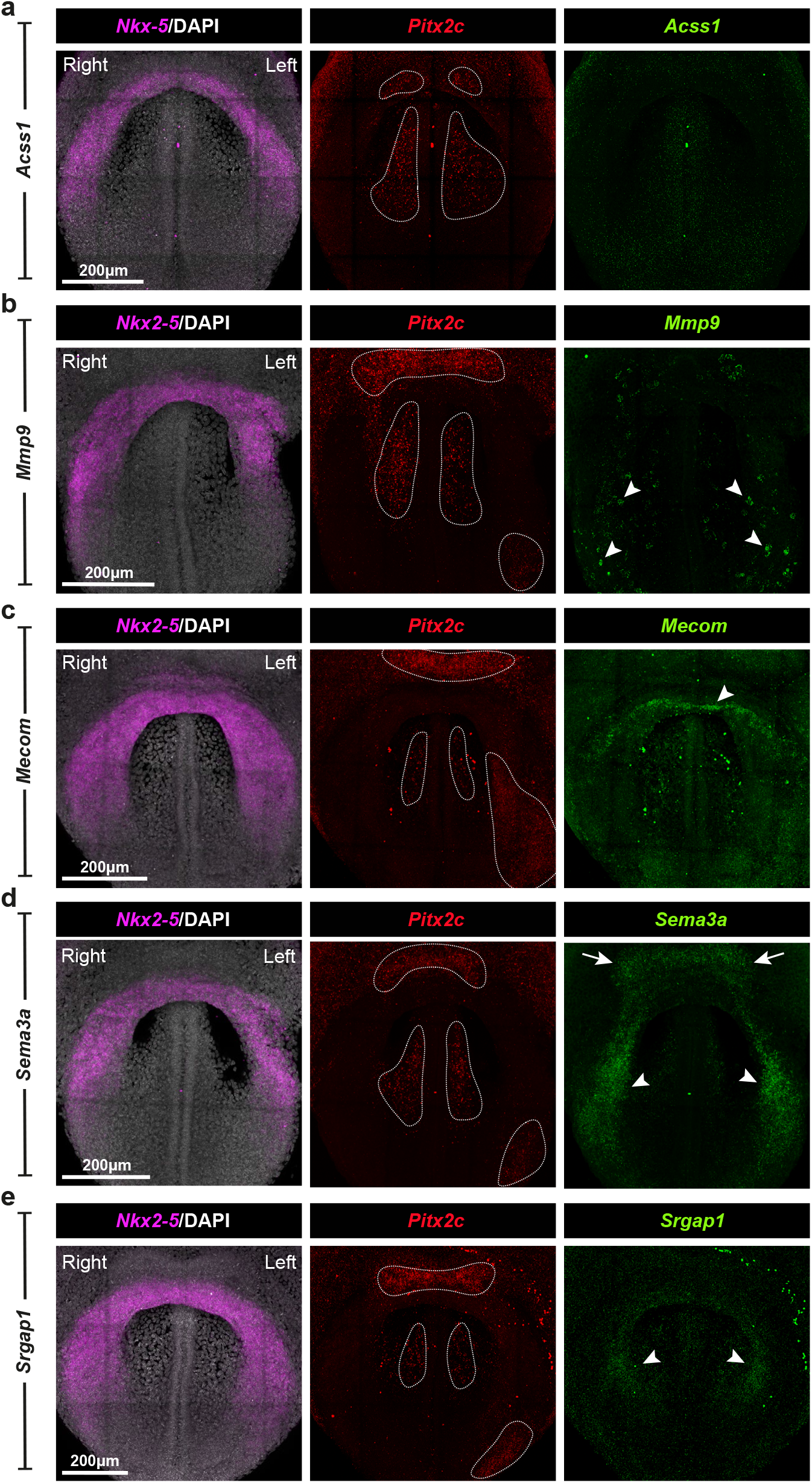
Expression of newly identified asymmetrically expressed genes during the onset of symmetry breaking. **a-e**, Maximum Intensity Projections (MIP) of hybridization chain reaction (HCR) staining revealing the expression pattern of *Acss1* (**a**), *Mmp9* (**b**), *Mecom* (**c**), *Sema3a* (**d**) and *Srgap1* (**e**), as well as *Pitx2c* and Nkx2-5, in embryos at the initiation of asymmetry breaking. Dotted regions highlight the symmetrical expression of *Pitx2c* in the ectoderm and paraxial mesoderm as well as expression in the left caudal lateral plate mesoderm (LPM). Arrowheads mark: individual *Mmp9* expressing cells (**b**); symmetrical Mecom expression in the endocardium (**c**); symmetrical *Sema3a* and *Srgap1* expression in LPM (**d** and **e**). Arrows mark *Sema3a* expression in the ectoderm (**d**).

Left-right asymmetric gene expression is triggered by events at the node dependent on leftward flow created by the action of nodal cilia^1,25^. Inversus viscerum (iv) mutant embryos have defective nodal cilia and randomized situs due to a mutation in *Dnah11*, the gene encoding the dynein axonemal heavy chain 11^26–28^. To test the dependence of the asymmetric expression of *Mecom* on nodal ciliary action, we looked at *Mecom* expression in *Dnah11*^*iv/iv*^ embryos. Since *Mecom* shows both symmetric (endocardium) and asymmetric (myocardial progenitors in LPM) expression within the forming heart, it offers a particularly interesting candidate to specifically study the asymmetric cardiac expression, using the symmetric component of expression as an internal control.

Homozygous *Dnah11*^*iv/iv*^ mutants are viable and have been described based on gross morphology to show a range of phenotypes ranging from *situs solitus* to *situs inversus totalis*^26,27^. We first performed a detailed molecular characterization of the left-right phenotype of these mutants during cardiac development stages, using *Pitx2c* expression as a readout of situs. In homozygous mutant embryos between stage 0 and LHT (n=90), *Pitx2c* showed four different expression profiles as previously reported^29^: left-asymmetric expression (32.22%), right-asymmetric expression (31.11%), symmetric expression (14.44%) and no expression (22.22%) (Figure 5a and 5c). During cardiac crescent to LHT maturation there was a shift in the proportion of embryos displaying a given *Pitx2c* expression profile (Figure 5b). In mutant embryos, the onset of asymmetric *Pitx2c* expression was delayed in comparison to that of wild type embryos, with asymmetric expression being detected only from stage 1 onwards. As the cardiac crescent transitioned into the LHT, the proportion of embryos with symmetrical *Pitx2c* expression increased, whilst the number of embryos with no *Pitx2c* expression decreased.

**Figure 5:**
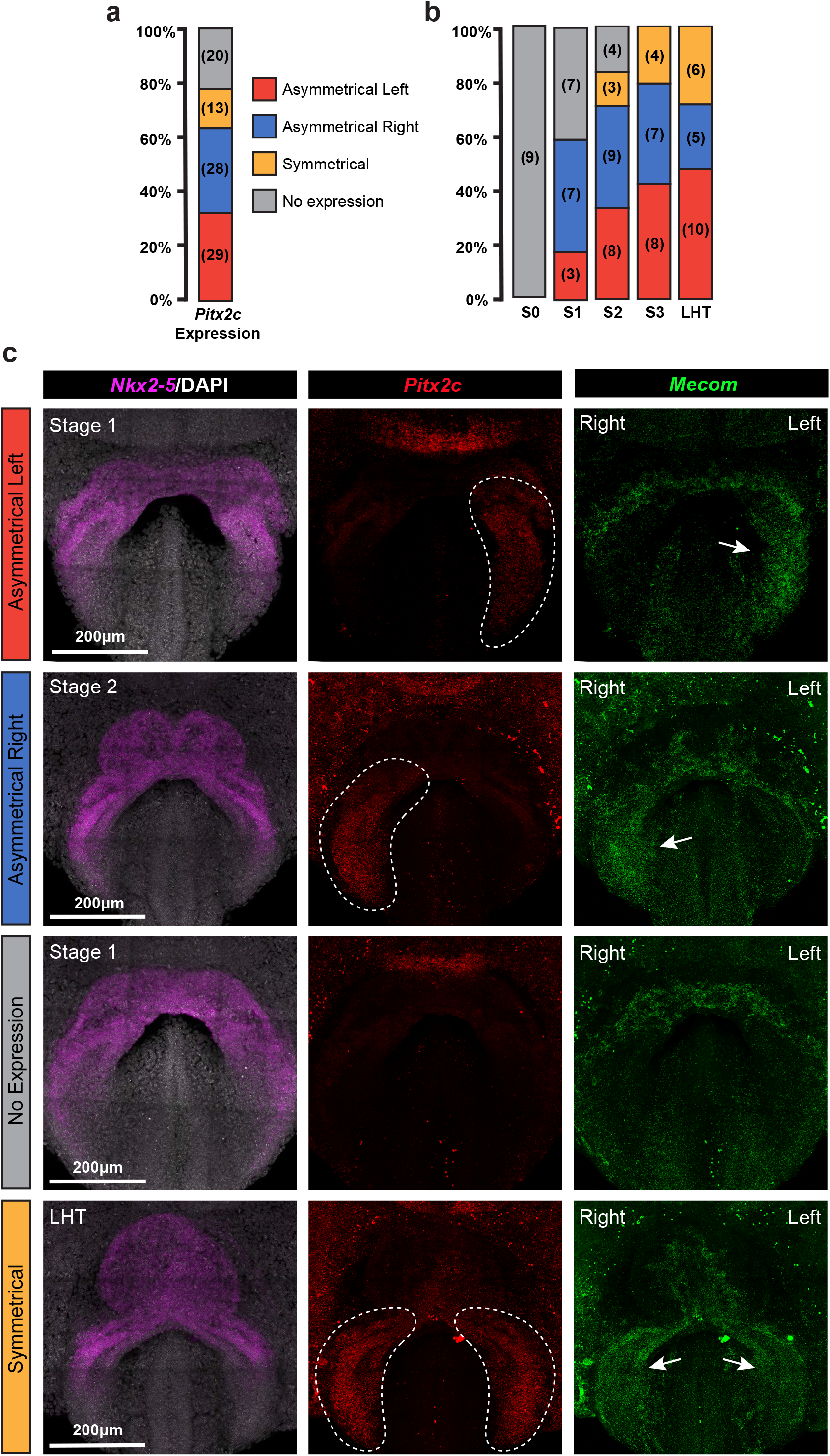
Characterization of Dnah11^iv/iv^ mutant phenotypes and Mecom expression. **a-b**, Combined and stage-specific quantification of *Pitx2c* expression patterns in iv/iv mutant embryos during cardiac crescent development. Number of embryos shown in brackets. **c**, Maximum Intensity Projections (MIP) of hybridization chain reaction (HCR) staining revealing the expression pattern of *Mecom*, in embryos with different *Pitx2c* expression profiles. Dotted regions represent *Pitx2c* expression and arrows mark *Mecom* expression.

When we looked at *Mecom* expression in these embryos, we observed normal symmetrical expression in the cardiac endothelium in all *iv/iv* mutant embryos irrespective of the expression pattern of *Pitx2c*. In mutant embryos in which *Pitx2c* was expressed asymmetrically in either the right or left LPM, we found that the LPM expression of *Mecom* correlated with the pattern of expression of *Pitx2c* (Figure 5c). In embryos where no *Pitx2c* expression was detected in the LPM, *Mecom* could only be detected in endothelial cells and was not expressed in LPM, whilst in embryos with symmetrical *Pitx2c* expression in the LPM, *Mecom* too was expressed symmetrically in the left and right LPM. Overall, these results indicate that *Mecom* expression in the endothelium and LPM are independently regulated and that the asymmetric LPM expression is determined by the same mechanisms that control asymmetric *Pitx2c* expression.

## DISCUSSION

Given the importance of left-right asymmetry in the formation of the body plan, it is fundamental to understand how these asymmetries arise during development. Our study characterizes the molecular differences which are present during symmetry breaking in the mouse embryo, specifically in the context of the embryonic heart, which is the first organ during development to manifest morphological indications of left-right asymmetry^18^.

Having previously mapped and characterized the cell types present in the anterior heart forming region of the embryo, we used this dataset to identify asymmetrically expressed genes and validated their expression pattern using HCR. Overall left-right transcriptional differences were subtle, having no influence on cell clustering. However, due to our manual sub-dissection, we retained information about whether cells originate on the left or right sides of the cardiac crescent and could leverage this information to identify genes that were expressed with a left or right sided bias. The vast majority of asymmetrically expressed genes were identified within mesodermal cell populations, highlighting germ layer specific responses to symmetry breaking at the node. In contrast to previous attempts at identifying left-right asymmetric gene expression, that identified genes relating predominantly to cardiomyocyte differentiation^20^, the asymmetric genes we have identified and validated play roles in a wide range of biological processes, highlighting the complexity by which asymmetry manifests. These include extracellular matrix remodeling (*Mmp9*), signaling (*Sema3a*), cell cycle progression (*Mecom*) and translation regulation (*Ybx3*). Apart from *Mmp9*, none of these genes have previously been implicated in left-right asymmetry breaking and therefore it would be important to further test and characterize the functional role of these genes in this process. Interestingly, homozygous Evi1 (*Mecom*) knockout mice die at around 10.5dpc. Mutant embryos had widespread hypocellularity and hemorrhaging, and hearts which had a more tube-like appearance, potentially suggesting a looping defect^30^.

Given our previous high-fidelity mapping of cell types in the cardiac region of the embryo^21^, we could also validate asymmetric gene expression biases within distinct anatomical locations. For example, *Mmp9* was most strongly expressed in cells from transcriptional clusters Me7, Me6, Me3 and endothelial cells (Me2). Anatomical validation of asymmetric *Mmp9* expression in the embryo showed expression in a specific region of the cardiac crescent which marked the transition from cardiac progenitors to cardiomyocytes (Figure S6b)^21^. This anatomical region transcriptionally represented cells from cluster Me6, which strongly expressed *Mmp9*, highlighting how this dataset can identify specific L/R asymmetries in transcriptionally and anatomically distinct populations of cells. Given the role of MMP9 in extracellular matrix remodeling it would suggest that this early asymmetric *Mmp9* expression plays a role in the onset of LHT looping.

Our computational analysis revealed several genes with right sided bias, which is striking given that relatively few right-asymmetric genes have been reported in the mouse embryo. Two such examples are *Cerl2*, whose transcripts are stabilized preferentially on the right side of the node; and *Snai1*, expressed more strongly in the right LPM than left^31–35^. When validating right-sided genes using HCR, we found a greater degree of variability in their expression patterns compared to left-sided genes. They did not always show right-sided expression, suggesting potential intrinsic differences in the regulation of right-sided genes compared to the left. This variability could reflect a less-stable, more transient, expression profile for right-sided gene expression relative to the left, which is robustly controlled by the node and downstream *Pitx2c* activation. Alternatively, it could reflect the overall high levels of expression of right sided genes in all cell types, resulting in a technical inability to detect relatively small or localized changes in gene expression levels.

One of the most significant right-sided genes was *Ybx3*, which encodes a DNA and RNA binding protein. YBX3 post-translationally controls mRNA stability and translation, having a role in the regulation of cell proliferation and density as well as in amino acid uptake^24,36^. Asymmetric proliferation rates have been reported to play a role in driving rightward looping of the LHT, with increased rates of proliferation of the right side of the venous pole^19,37^. Reduction of YBX3 in vitro leads to a decrease in proliferation, potentially highlighting that it may play a role in regulating asymmetric proliferation differences during early cardiac morphogenesis.

We have observed the spread of *Pitx2c* expression, from a distal region of LPM adjacent to the node, proximally through the LPM and into the cardiac crescent. In contrast we have observed the maintenance of symmetric *Pitx2c* expression in the head folds and yolk sac endoderm, highlighting tissue specific regulation of this gene. Interestingly, we discovered other genes which showed tissue specific asymmetries, such as *Mecom*. Prior to the asymmetric expression of *Pitx2c, Mecom* could be detected symmetrically in the endothelial cells of the presumptive endocardium. However, after *Pitx2c* became asymmetric, *Mecom* too became asymmetrically expressed in the LPM, while remaining symmetrically expressed in the endocardium. This highlights differences in the regulation of asymmetric genes within similar anatomical regions but in a cell type specific manner (i.e. endothelial vs. LPM) and could reflect how asymmetric signals propagate from the node but also how these different cell types respond to the asymmetry generating signals. It also indicates that within the heart, different cell types contribute disproportionately to the generation of morphological asymmetries.

This study represents a comprehensive transcriptional characterization of asymmetry breaking during cardiac crescent development, in the framework of a single-cell dataset. Given the central role left-right asymmetry plays in cardiac morphogenesis, it provides a valuable resource for further mechanistic studies, aiding in our understanding of congenital heart diseases.

## MATERIALS AND METHODS

### Mouse strains, husbandry and embryo collection

All animal experiments complied with the UK Animals (Scientific Procedures) Act 1986, approved by the local Biological Services Ethical Review Process and were performed under UK Home Office project licenses PPL 30/3420 and PCB8EF1B4. To obtain wild-type embryos, C57BL/6 males (in house) were crossed with 8-16 week old CD1 females (Charles River, England). All mice were maintained in a 12-hr light-dark cycle. Noon of the day when a vaginal plug was found was designated E0.5. To dissect the embryos, the pregnant females were culled by cervical dislocation in accordance with schedule one of the Animals (Scientific Procedures) Act. Embryos of the appropriate stage were dissected in M2 medium (Sigma-Aldrich, Cat No. M7167).

### Dissection and unbiased collection of cells from the mouse cardiac crescent

Following dissection, embryos were collected, keeping the yolk sac intact, in fresh M2 media before being grouped based on their cardiac crescent stage. Progressive crescent stages (−1 to 3) were defined based on morphological criteria including the shape of the cardiac crescent and the ratio between width (medio-lateral axis) and maximum height (rostral-caudal axis) of the crescent^38^. Stage -1 embryos were defined based on the incomplete fusion of the mesodermal wings in the anterior portion of the embryo. Embryos were considered to be at the LHT stage once both sides of the cardiac crescent were completely folded and fused. Embryos prior to cardiac crescent formation were staged using the Lawson and Wilson staging system of mouse development^39^.

Once embryos were staged, crescent regions were micro-dissected using tungsten needles, keeping the overlying endoderm intact. By maintaining the integrity of the overlying endoderm at cardiac crescent and LHT stages, regions of both yolk sac endoderm and embryonic endoderm were also collected as well as the forming pericardium. To isolate cardiac crescent regions, the anterior half of the embryo was dissected from the posterior before a cut was made along the midline separating the embryo into left and right sides. The cardiac crescent could then be isolated using the sagittal profile to collect both dorsal and ventral tissue. Left and right sides of the sub-dissected regions where disaggregated and processed separately. At LHT stage the heart tube was isolated by dissections from the ventral surface, due to both the morphology and size of the tissue, therefore it is unlikely we collected regions of the dorsal pericardial wall. Following isolation of the cardiac crescent region the tissue was pooled from multiple embryos (Embryos pooled: stage LHT: 5, stage 3; 7, stage 2; 27, stage 1; 18, stage 0; 7, stage -1; 2). The pooled samples were dissociated into single cells using 200μl Accutase (ThermoFisher, Cat No. A1110501) for 12 minutes at 37°C, being agitated every 2 minutes, before adding 200μl heat-inactivated FBS (ThermoFisher, Cat No. 10500) to quench the reaction. Cells were then centrifuged at 1000rpm for 3 minutes at 4°C before being suspended in 100μl HBSS (ThermoFisher, Cat No. 14025) + 1% FBS, and stored on ice. Single cells were collected using a Sony SH800 FACS machine with a stringent single-cell collection protocol and sorted into 384 well plates containing SMART-seq2 lysis buffer^40^ plus ERCC spike-ins. To ensure we collected good quality cells, a live/dead dye (Abcam, Cat No. ab115347) was used; 100μl was added to the cell suspension at a 2x concentration in HBSS 10 minutes before collection, and live cells were collected based on their FITC intensity. Once cells were collected, plates were sealed, spun down, and frozen using dry ice before being stored at -80°C. This complete process, from dissection to single-cell collection, took approximately 2-3 hours. Cells were collected in multiple batches.

### Differential expression analysis

To identify asymmetrically expressed genes, we used the previously reported data from Tyser et al, 2021, obtained from https://content.cruk.cam.ac.uk/jmlab/mouseEmbryonicHeartAtlas/. Briefly, single-cell transcriptomic data for good-quality cells is available in a SingleCellExperiment object (sce_goodQual.NORM.clusters.Rds), containing raw and normalized counts, together with batch, stage, side, and cluster annotations for each cell. This object was the basis for all analyses reported in this study.

To test for differential expression between cells from the left and right sides, we converted the SingleCellExperiment object into a DGElist object (convertTo function from scran package (Lun, McCarthy, Marioni, F1000Res, 2016)) to be used with edgeR (Robinson, McCarthy, Smyth, Bioinformatics, 2010). We analyzed each germ layer separately and used generalized linear models to test for differences between left and right sides in cells from each cluster (with glmQLFit and glmQLFTest), while controlling for batch (experiment date). Contrasts for all clusters from the same germ layer were tested together to reduce the number of tests performed; we thus obtained a single p-value indicating whether the gene is differentially expressed in *any* of the clusters tested (Me1 and Me2 were excluded due to their very small number of cells). P-values were adjusted for multiple testing using the Benjamini Hochberg method.

To define significant changes in expression, we required a gene to have an adjusted p-value smaller than 0.05 and an absolute fold-change larger than 1.5 for mesoderm clusters, and 1.3 for endoderm and ectoderm clusters. We filtered out genes with unstable fold-changes that stem from outlier expression in a small number of cells. For this, we computed the average gene expression levels in each cluster and ignored fold-change estimates from clusters where gene expression was lower than 1 normalized log_2_ count. We filtered out any genes that no longer fulfilled the minimum fold-change threshold from our list of significant results. This procedure ensures that the asymmetrically expressed genes defined are supported by at least moderate expression levels in the clusters showing differential expression. Full differential expression results are provided in Supplementary Table 2; fold-change estimates for clusters with very low expression are set to NA.

### Cell-cycle phase differences

To test whether there are significant differences in the proportions of cells in non-cycling (G1) versus cycling (S and G2/M) cells between the left and right sides, we used a binomial test (with the proportions on the left used as the expected probabilities, and testing whether the proportions on the right were significantly different). Cell-cycle phase assignment for each cell was originally inferred using Cyclone (Scialdone et al., 2015).

### Whole mount immunostaining

Dissected intact embryos were fixed for 1 hr at room temperature with 4% paraformaldehyde (SantaCruz, Cat No. sc281692) in PBS (Sigma-Aldrich, Cat No. P4417). The embryos were then rinsed twice in PBST-0.1% (PBS with 0.1% Triton X-100 (Sigma-Aldrich, Cat No. T8787)) before being permeabilized in PBST-0.25% for 40 mins and rinsed again 3x in PBT-0.1%. The embryos were next transferred to blocking solution (5% donkey serum (Sigma-Aldrich, Cat No. D9663), 1%BSA (Sigma-Aldrich, Cat No. A7906) in PBST-0.1%) overnight (o/n) at 4°C. Primary antibodies (Goat Nkx2-5, Santa Cruz) were then added to the solution and were incubated o/n at 4°C. The embryos were washed 3x 15mins in PBST-0.1% and incubated o/n at 4°C in PBST-0.1% with the secondary antibodies, and DAPI, then subsequently washed 3x PBT-0.1% for 15 min and placed in Vectashield mounting medium with DAPI (Vector Labs, Cat No. H-1200) for at least 24 hr at 4°C. Each staining combination was repeated on at least 3 embryos. Samples were imaged using a Zeiss 880 confocal microscope with a 40x oil (1.36 NA) objective. Images were captured at 512 × 512 pixel dimension using multiple tiles with a Z-step of between 1.5 and 0.5 µm. Based on the working distance of the Zeiss 880 microscope and tissue penetration, embryos were imaged to a depth of between 120 – 150µm.

### In Situ Hybridization Chain reaction (HCR)

In situ HCR kit (ver.3) containing amplifier set, hybridization, amplification, wash buffers, and DNA probe sets, were purchased from Molecular Instruments (molecularinstruments.org) and the protocol described in Choi et al. (2018) was followed with slight modifications^41^. Embryos were collected as previously described and fixed overnight in 4% paraformaldehyde at 4°C with shaking. Following fixation, the embryos were kept on ice and washed twice in cold nuclease free 0.1% PBS Tween (nucPBST) before being dehydrated in methanol with a series of graded MeOH/0.1% nucPBST washes (25% MeOH/75% nucPBST; 50% MeOH/50% nucPBST; 75% MeOH/25% nucPBST; 2x 100% MeOH) for 10 minutes on ice before being stored at -20°c in 100% MeOH. For HCR staining, embryos were rehydrated with a series of graded MeOH/PBST washes for 10 minutes each on ice (75% MeOH/25% nucPBST; 50% MeOH/50% nucPBST; 25% MeOH/75% nucfree PBST; 100% nucPBST; 100% nucPBST at room temperature). After rehydration, embryos were treated with 10 μg/ml proteinase K solution for 5 minutes at room temperature, washed twice for 5 minutes with nucPBST, post-fixed with 4% paraformaldehyde for 20 minutes at room temperature, then washed three times with nucPBST for 5 minutes. Embryos were next pre-hybridized with 50% probe hybridization buffer/ 50% nucPBST at room temperature for two hours before being incubated with probe hybridization buffer for 30 minutes at 37°C prior to being incubated in probe solution (2 pmol of each probe in 500μl of 30% probe hybridization buffer) overnight (12-16 hours) at 37°C. Probe libraries were designed and manufactured by Molecular Instruments using *Mus musculus* sequences from NCBI database. Excess probe was removed by washing with pre-warmed probe wash buffer at 37°C four times for 15 minutes each and then two times 5 minutes with 5X SSCT buffer (5x sodium chloride sodium citrate, 0.1% Tween 20 in ultrapure H2O). For amplification, embryos were incubated in amplification buffer for 30 minutes at room temperature before being placed into hairpin solution and incubated overnight in the dark at room temperature. 30 pmol hairpin solution was prepared by snap cooling (95°C for 90 seconds, then cooled to room temperature) stock hairpins in storage buffer and then diluted in amplification buffer at room temperature. Following overnight incubation, excess hairpins were removed by washing with 5X SSCT buffer at room temperature (2x 5 mins, 2x 30 mins with DAPI and 1x 5 min). Embryos were then placed into Vectashield antifade mounting medium (Vector Laboratories, Cat no. H-1000) or 87% glycerol solution, then imaged on a Zeiss 880 confocal microscope as previously described in the immunohistochemistry section. Each HCR combination was repeated on at least 3 embryos. Sagittal sections were generated using optical reconstruction of multiple Z-images.

## FUNDING

This work was funded by Wellcome Awards 108438/Z15 and 105031/C/14/Z (SS and JCM), Wellcome Senior Investigator Award 103788/Z/14/Z (SS), BHF Immediate Fellowship FS/18/24/33424 (RT) and core funding from the European Molecular Biology Laboratory and Cancer Research UK (JCM).

## AUTHOR CONTRIBUTIONS

*in situ* HCR imaging: RT, MP; computational analyses: XI-S; whole mount immunohistochemistry: RT; dissection and processing embryos for single-cell library preparation RT, SS and AM; initial scRNA-seq data analyses: TvdB and AS; Preparation of manuscript draft: RT; Preparation of figures: RT, XI-S; Editing and review of final manuscript: RT, XI-S, MP, AM, TvdB, AS, JM, SS. JM and SS supervised the study. All authors read and approved the final manuscript. Authors declare no competing interests.

## DATA AND MATERIALS AVAILABILITY

This study utilizes previously reported data (Tyser, Ibarra-Soria et al, Science, 2021), obtained from https://content.cruk.cam.ac.uk/jmlab/mouseEmbryonicHeartAtlas/. All code used for data analysis and figure generation is available at https://github.com/MarioniLab/leftRight_asymmetry.

**Figure S1:**
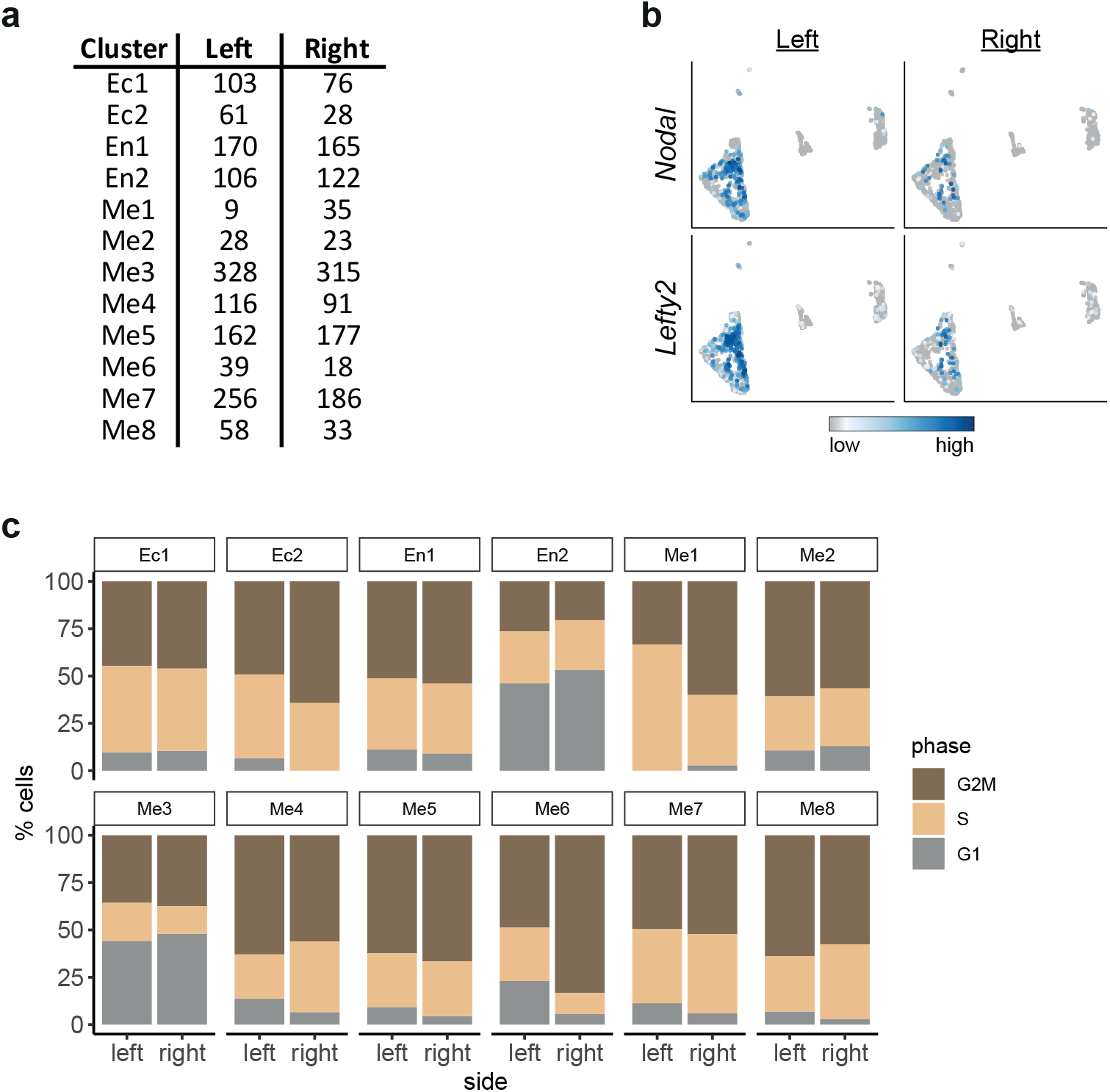
Characterization of left right differences during cardiac crescent development. **a**, Table showing the number of cells from each transcriptional cluster collected from either the left or right side of the embryo. **b**, UMAP showing the expression of Lefty2 and Nodal in cells collected from either the left or right side of the forming cardiac crescent. **c**, Bar graphs showing the proportion of left and right cells from different cell clusters in either G1, S or G2M phase of the cell cycle.

**Figure S2:**
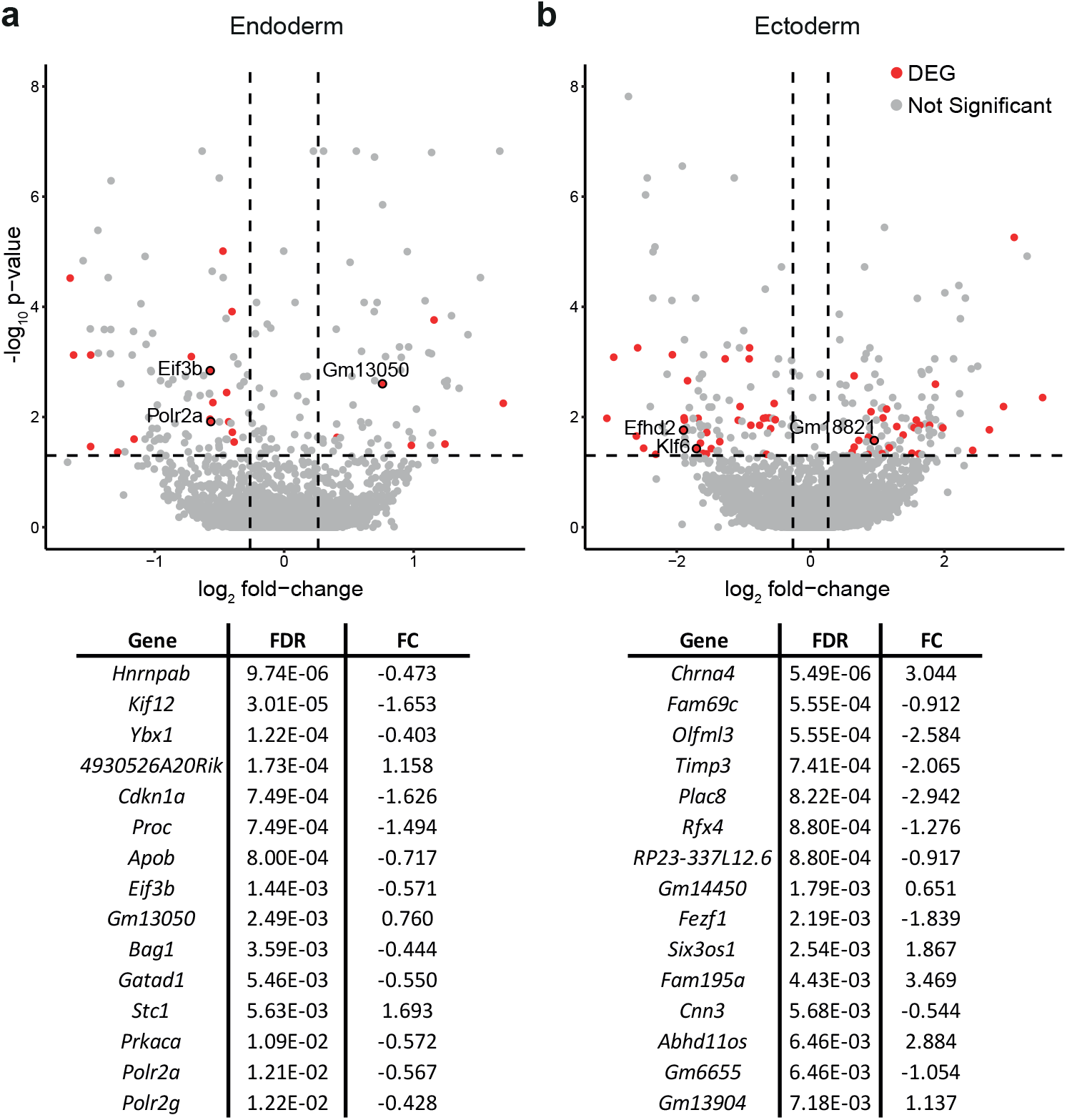
Identification of asymmetrically expressed genes in the endoderm and ectoderm. Differential expression analysis in the endodermal (**a**) and ectodermal (**b**) cells. The adjusted p-value (y-axis) is plotted against the fold-change between left-right cells (x-axis). For non-differentially expressed genes (grey), the average fold-change for all clusters is used; for DEGs (red) the largest fold-change (in clusters where the gene is expressed) is used. Genes are considered DE if their adjusted p-value is lower than 0.05 and their fold-change is larger than 1.3 (indicated by the dashed lines). Tables below show the top 15 DEGs, along with false discovery rate (FDR) and log fold change (FC) when comparing cells collected from either the left or the right.

**Figure S3:**
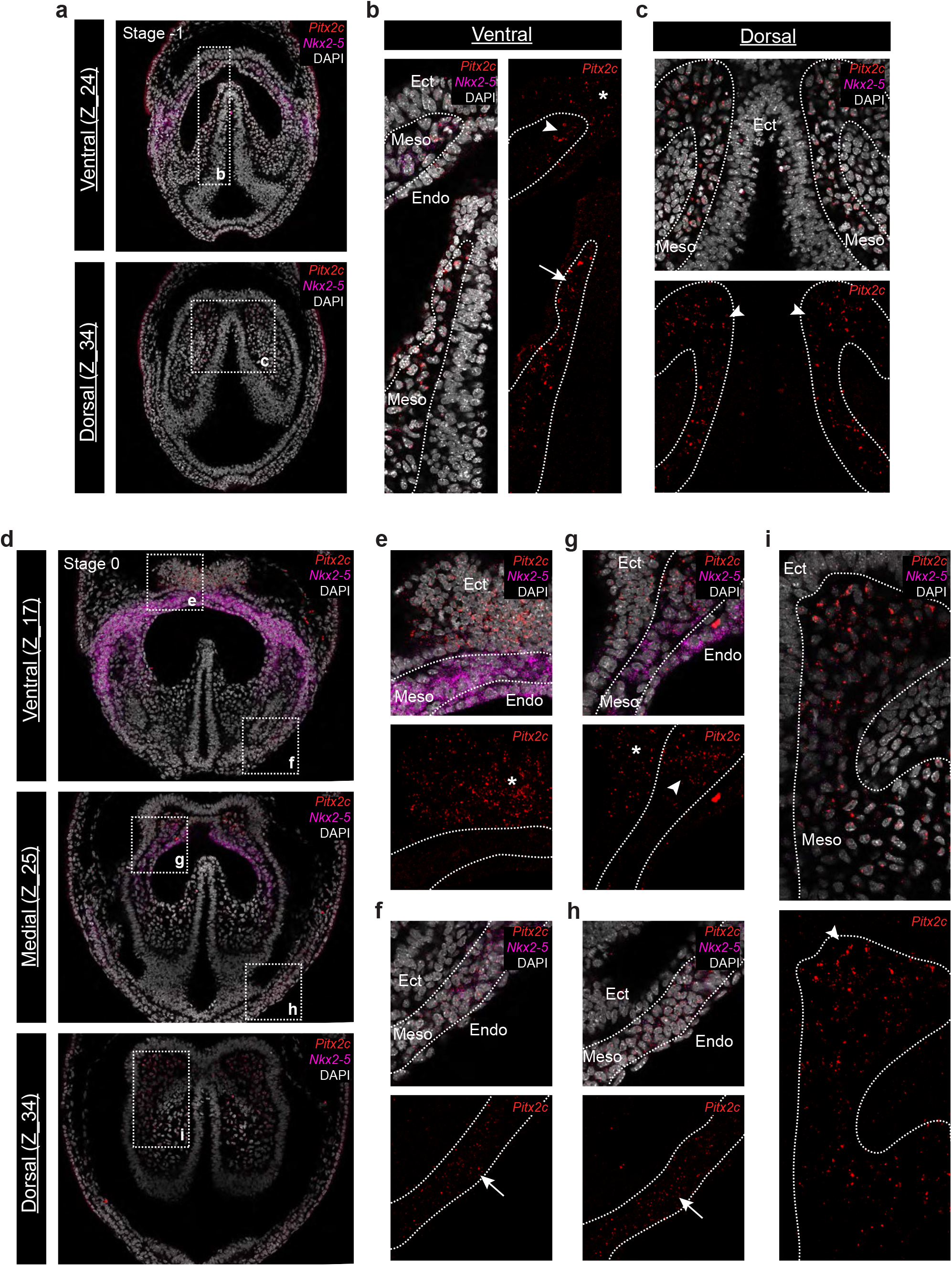
Detailed tissue specific characterization of Pitx2c expression during the onset of symmetry breaking. **a**, Individual ventral (top) and dorsal (bottom) sections of hybridization chain reaction (HCR) staining revealing the expression pattern of *Pitx2c* and *Nkx2-5* in a stage -1 cardiac crescent embryo (Figure 2a). **b**, Insert from panel a showing the ventral expression of *Pitx2c* in the paraxial (arrow) and cranial (arrowhead) mesoderm and neurectoderm (asterisks). Meso; mesoderm; Ecto, neuroectoderm; Endo, endoderm. **c**, Insert from panel a showing the dorsal expression of *Pitx2c* in the cranial mesoderm (arrowheads). **d**, Individual ventral (top), medial (middle) and dorsal (bottom) sections of hybridization chain reaction (HCR) staining revealing the expression pattern of *Pitx2c* and *Nkx2-5* in a stage 0 cardiac crescent embryo (Figure 2a). **e**, Insert from panel d showing the ventral expression of *Pitx2c* in the neurectoderm (asterisks). **f**, Insert from panel d showing the ventral expression of *Pitx2c* in the left distal lateral plate mesoderm (arrow). **g**, Insert from panel d showing the medial expression of *Pitx2c* in the cranial mesoderm (arrowheads) and neurectoderm (asterisks). **h**, Insert from panel d showing the medial expression of *Pitx2c* in the left distal lateral plate mesoderm (arrow). **i**, Insert from panel d showing the expression of *Pitx2c* in the dorsal cranial mesoderm (arrowhead).

**Figure S4:**
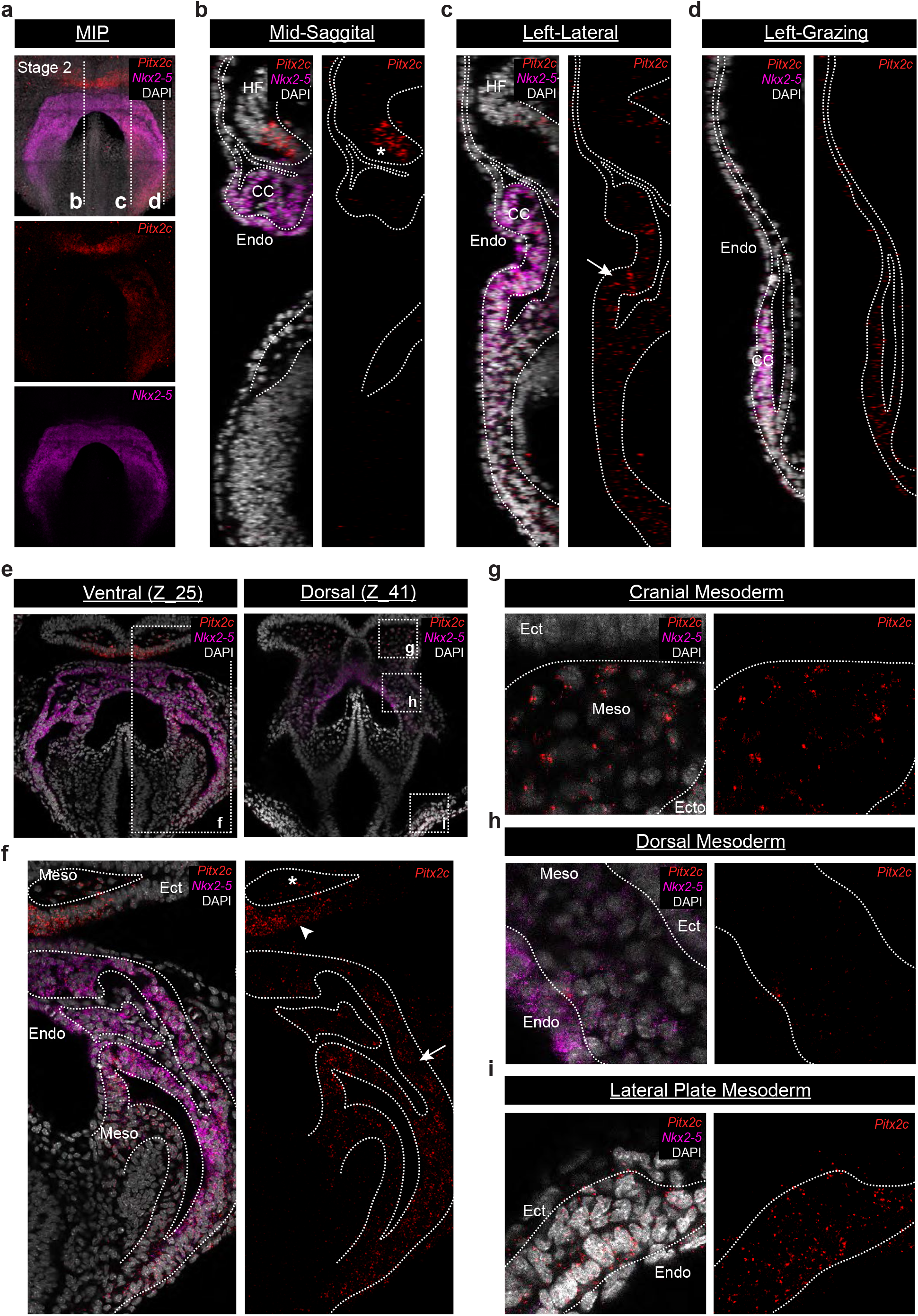
Expression pattern of Pitx2c in a stage 2 cardiac crescent. **a**, Maximum intensity projection (MIP) of a stage 2 cardiac crescent embryo showing expression of *Nkx2-5* and *Pitx2c* using Hybridization Chain Reaction (HCR). **b**, Mid-sagittal section of the embryo in panel a, showing expression of *Pitx2c* in neurectoderm (asterisks) and absence from the cardiac crescent (CC). Endo, endoderm; HF, headfold. **c**, Left-lateral section of the embryo in panel a, showing expression of *Pitx2c* in left lateral plate mesoderm and extending into the cardiac crescent (arrow). **d**, Left-grazing section of the embryo in panel a, showing expression of *Pitx2c* in distal left lateral plate mesoderm. **e**, Individual ventral (left) and dorsal (right) coronal sections of HCR staining from shown in figure S4a. **f**, Insert from panel e showing the ventral expression of *Pitx2c* in the left lateral plate mesoderm (arrows), cranial mesoderm (arrowheads) and neurectoderm (asterisks). Ect, ectoderm. **g, h, i**, Inserts from panel e showing the dorsal expresvsion pattern of *Pitx2c* in different left-sided mesoderm regions.

**Figure S5:**
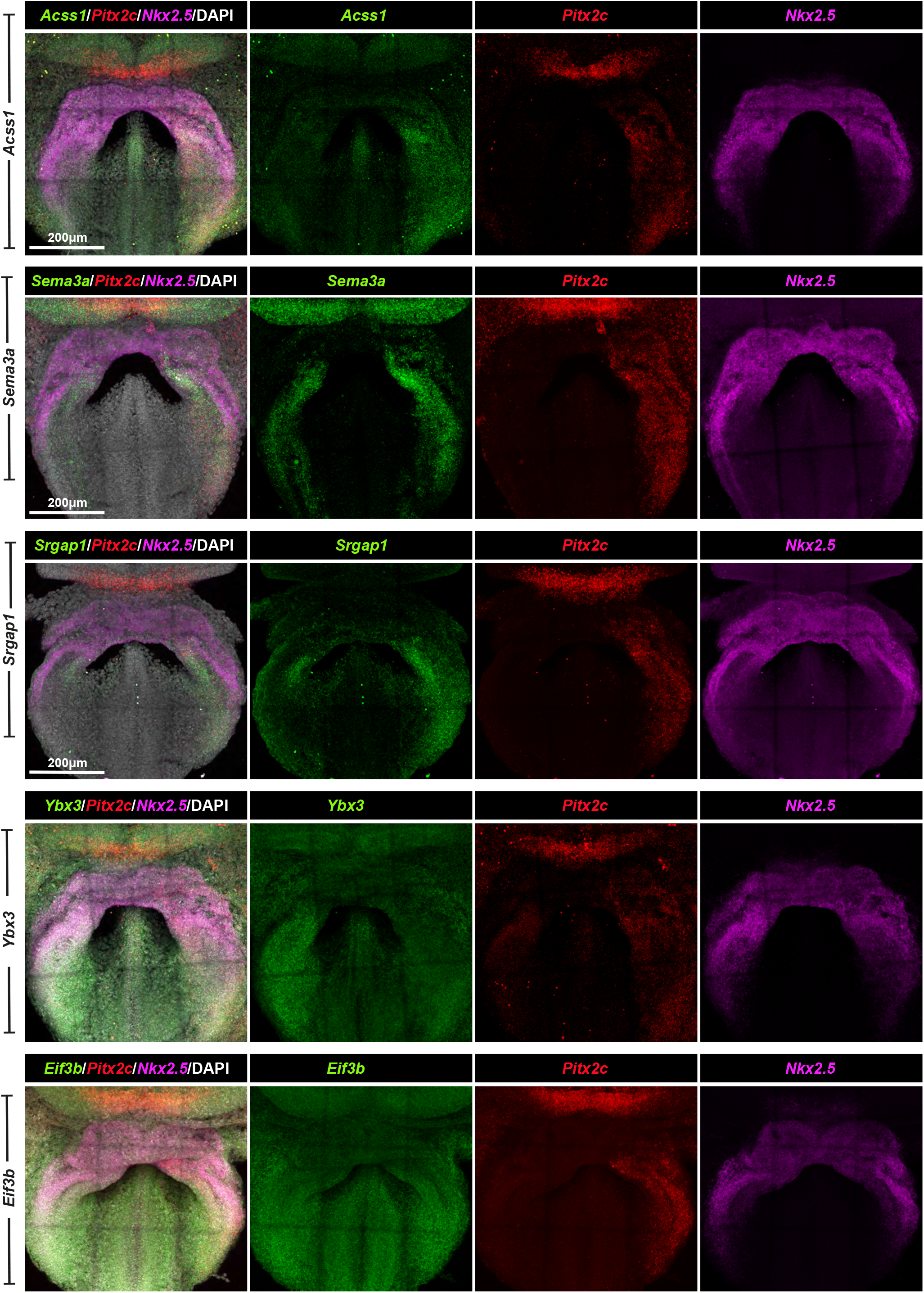
Individual channels for images in Figure 2. Maximum intensity projection (MIP) of a stage 2 cardiac crescent embryo showing expression of *Nkx2-5* and *Pitx2c* using Maximum Intensity Projections (MIP) showing merged and individual channels of hybridization chain reaction (HCR) staining from Figure 2c’-f’.

**Figure S6:**
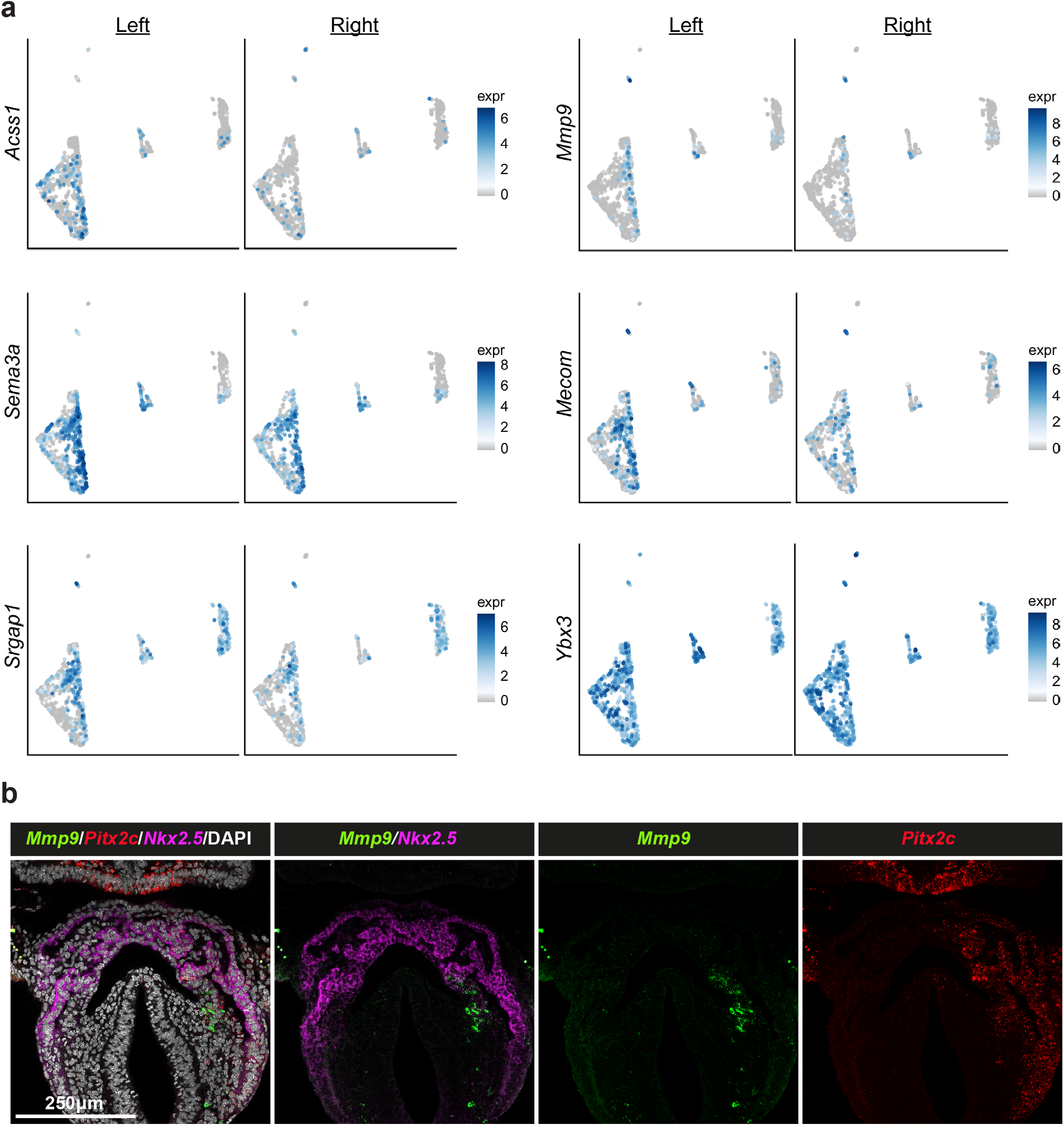
Asymmetric Mmp9 mesoderm is located in a distinct anatomical region of the developing cardiac crescent. a, UMAP showing the expression of *Acss1, Mmp9, Mecom, Srgap, Sema3a*, and *Ybx3* in cells collected from either the left or right side of the forming cardiac crescent. b, Individual sections from Figure 3e, showing the merged and separate expression pattern of *Mmp9, Pitx2c* and *Nkx2-5*.

**Figure S7:**
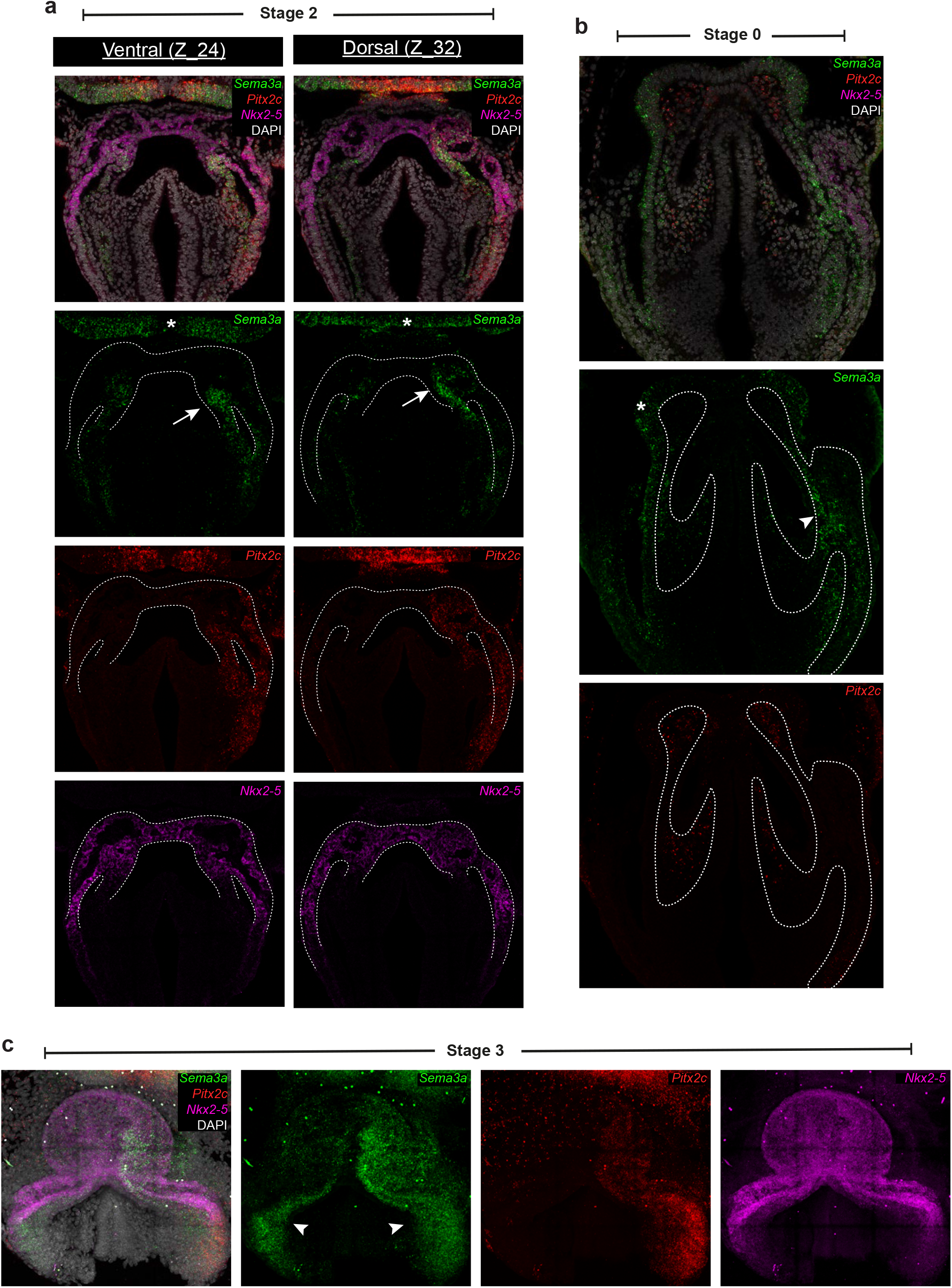
Expression of Sema3a in lateral plate mesoderm and neurectoderm. **a**, Ventral (left) and dorsal (right) Z sections showing hybridization chain reaction (HCR) staining for *Sema3a, Pitx2c* and *Nkx2-5* in a stage 2 cardiac crescent. Sema3a can be detected in both the neurectoderm (asterisks) as well as lateral plate mesoderm extending into the cardiac crescent (arrows). Dotted lines outline the cardiac crescent region. **b**, Z sections from Figure 4d showing hybridization chain reaction (HCR) staining for *Sema3a, Pitx2c* and *Nkx2-5* in a stage 0 cardiac crescent. *Sema3a* can be detected in both the neurectoderm (asterisks) as well as lateral plate mesoderm (arrows). Dotted lines outline mesoderm. **c**, Maximum intensity projection (MIP) of a linear heart tube stage embryo showing expression of *Sema3a, Nkx2-5* and *Pitx2c* using HCR. *Sema3a* expression extends throughout the left side of the heart tube but is also expressed in both venous poles (arrowheads).

**Supplementary Table 1:** Number of left/right cells detected in different cell-cycle phases.

| Left |  |  |  |  |  | Right |  |  |  |  |  |  |  |
| --- | --- | --- | --- | --- | --- | --- | --- | --- | --- | --- | --- | --- | --- |
| Cluster | Cell Number |  |  | Percentage |  |  | Cluster | Cell Number |  |  | Percentage |  |  |
|  | G1 | G2M | S | G1 | G2M | S |  | G1 | G2M | S | G1 | G2M | S |
| Ec1 | 10 | 46 | 47 | 9.7 | 44.7 | 45.6 | Ec1 | 8 | 35 | 33 | 10.5 | 46.1 | 43.4 |
| Ec2 | 4 | 30 | 27 | 6.6 | 49.2 | 44.3 | Ec2 | 0 | 18 | 10 | 0.0 | 64.3 | 35.7 |
| En1 | 19 | 87 | 64 | 11.2 | 51.2 | 37.6 | En1 | 15 | 89 | 61 | 9.1 | 53.9 | 37.0 |
| En2 | 49 | 28 | 29 | 46.2 | 26.4 | 27.4 | En2 | 65 | 25 | 32 | 53.3 | 20.5 | 26.2 |
| Me1 | 0 | 3 | 6 | 0.0 | 33.3 | 66.7 | Me1 | 1 | 21 | 13 | 2.9 | 60.0 | 37.1 |
| Me2 | 3 | 17 | 8 | 10.7 | 60.7 | 28.6 | Me2 | 3 | 13 | 7 | 13.0 | 56.5 | 30.4 |
| Me3 | 145 | 117 | 66 | 44.2 | 35.7 | 20.1 | Me3 | 151 | 118 | 46 | 47.9 | 37.5 | 14.6 |
| Me4 | 16 | 73 | 27 | 13.8 | 62.9 | 23.3 | Me4 | 6 | 51 | 34 | 6.6 | 56.0 | 37.4 |
| Me5 | 15 | 101 | 46 | 9.3 | 62.3 | 28.4 | Me5 | 8 | 118 | 51 | 4.5 | 66.7 | 28.8 |
| Me6 | 9 | 19 | 11 | 23.1 | 48.7 | 28.2 | Me6 | 1 | 15 | 2 | 5.6 | 83.3 | 11.1 |
| Me7 | 29 | 127 | 100 | 11.3 | 49.6 | 39.1 | Me7 | 11 | 97 | 78 | 5.9 | 52.2 | 41.9 |
| Me8 | 4 | 37 | 17 | 6.9 | 63.8 | 29.3 | Me8 | 1 | 19 | 13 | 3.0 | 57.6 | 39.4 |

**Supplementary Table 2a:**
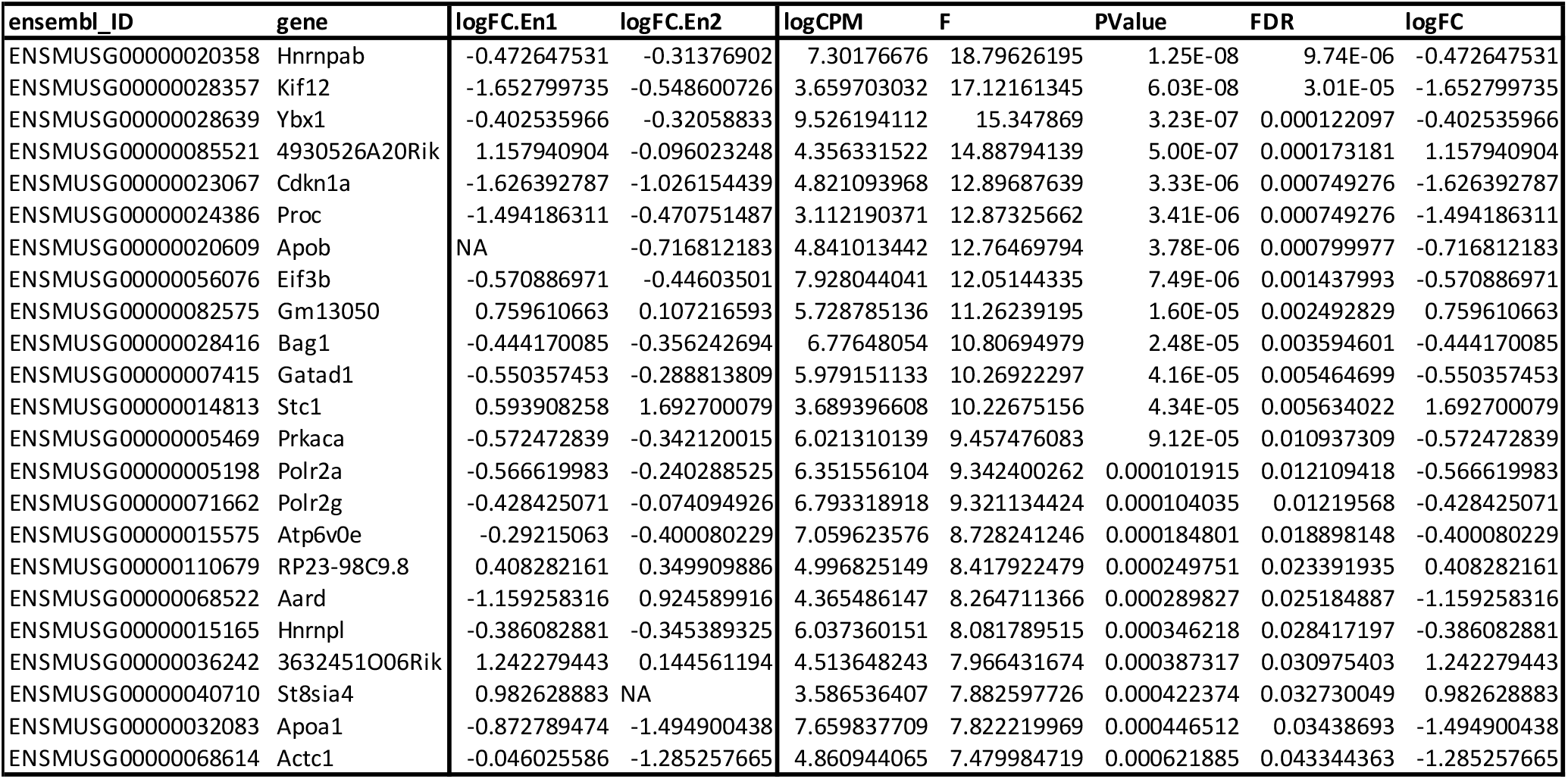
List of differentially expressed genes in left and right anterior endoderm. Results reported for all genes considered significantly differentially expressed. The fold-change in each cluster is reported. Clusters with very low expression were filtered out and the fold-change is NA. A summary fold-change is reported (**logFC**) and corresponds to the largest (absolute) fold-change from all clusters. Genes are considered significantly DE if FDR < 0.05 **AND** they show an absolute fold-change for at least one cluster larger than 1.3.

**Supplementary Table 2b:**
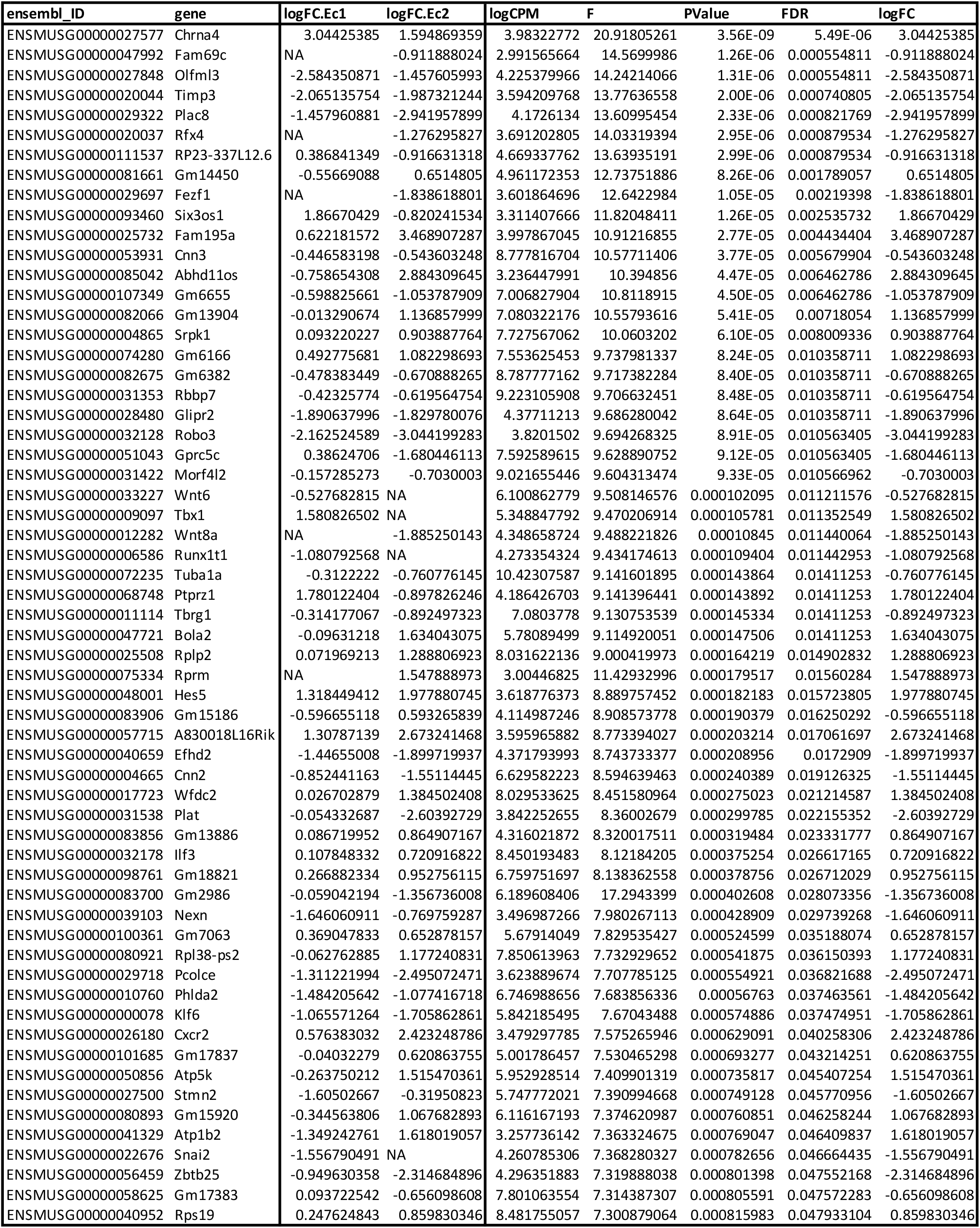
List of differentially expressed genes in left and right anterior ectoderm. Results reported for all genes considered significantly differentially expressed. The fold-change in each cluster is reported. Clusters with very low expression were filtered out and the fold-change is NA. A summary fold-change is reported (**logFC**) and corresponds to the largest (absolute) fold-change from all clusters. Genes are considered significantly DE if FDR < 0.05 **AND** they show an absolute fold-change for at least one cluster larger than 1.3.

**Supplementary Table 2c:**
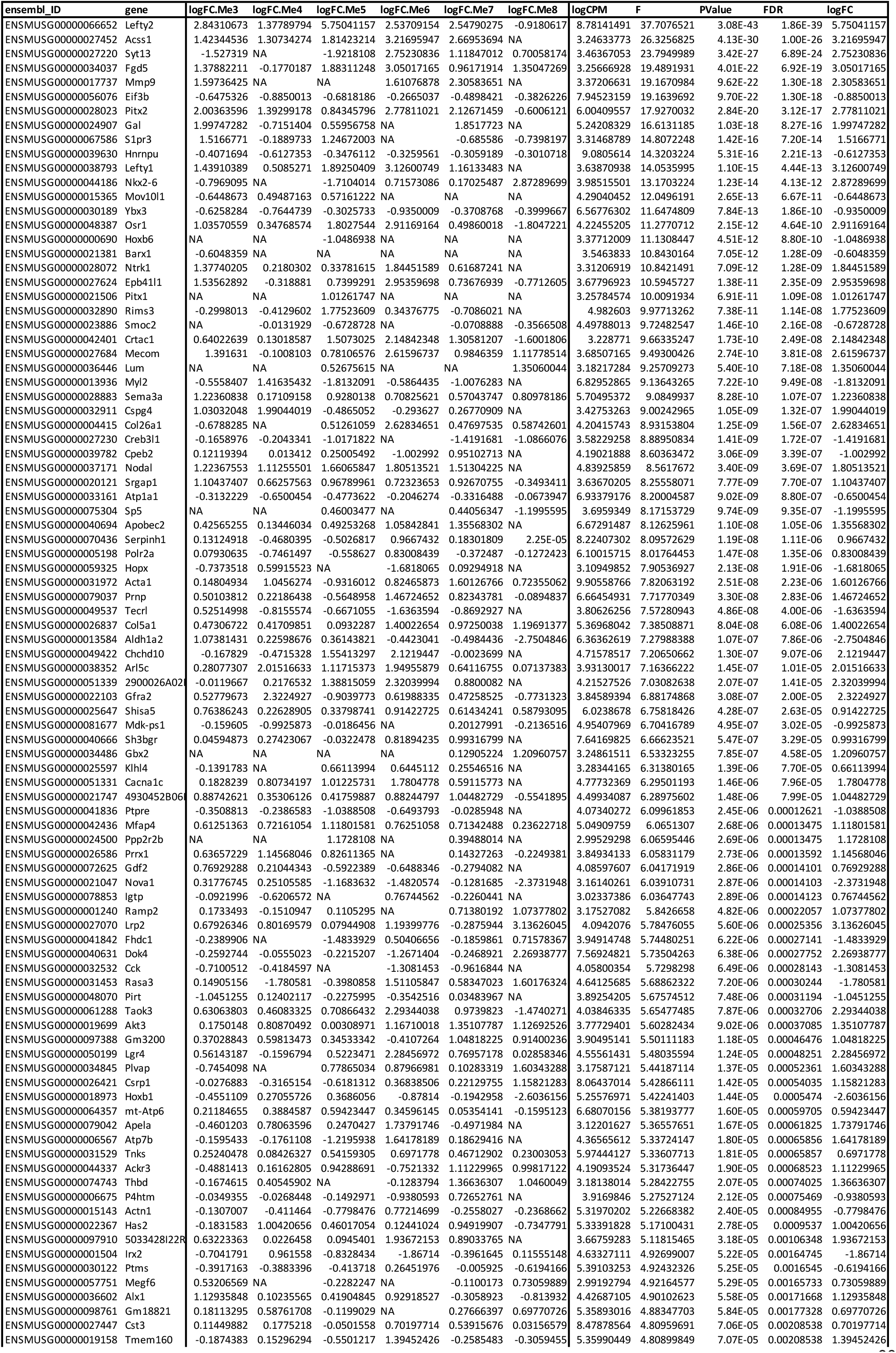
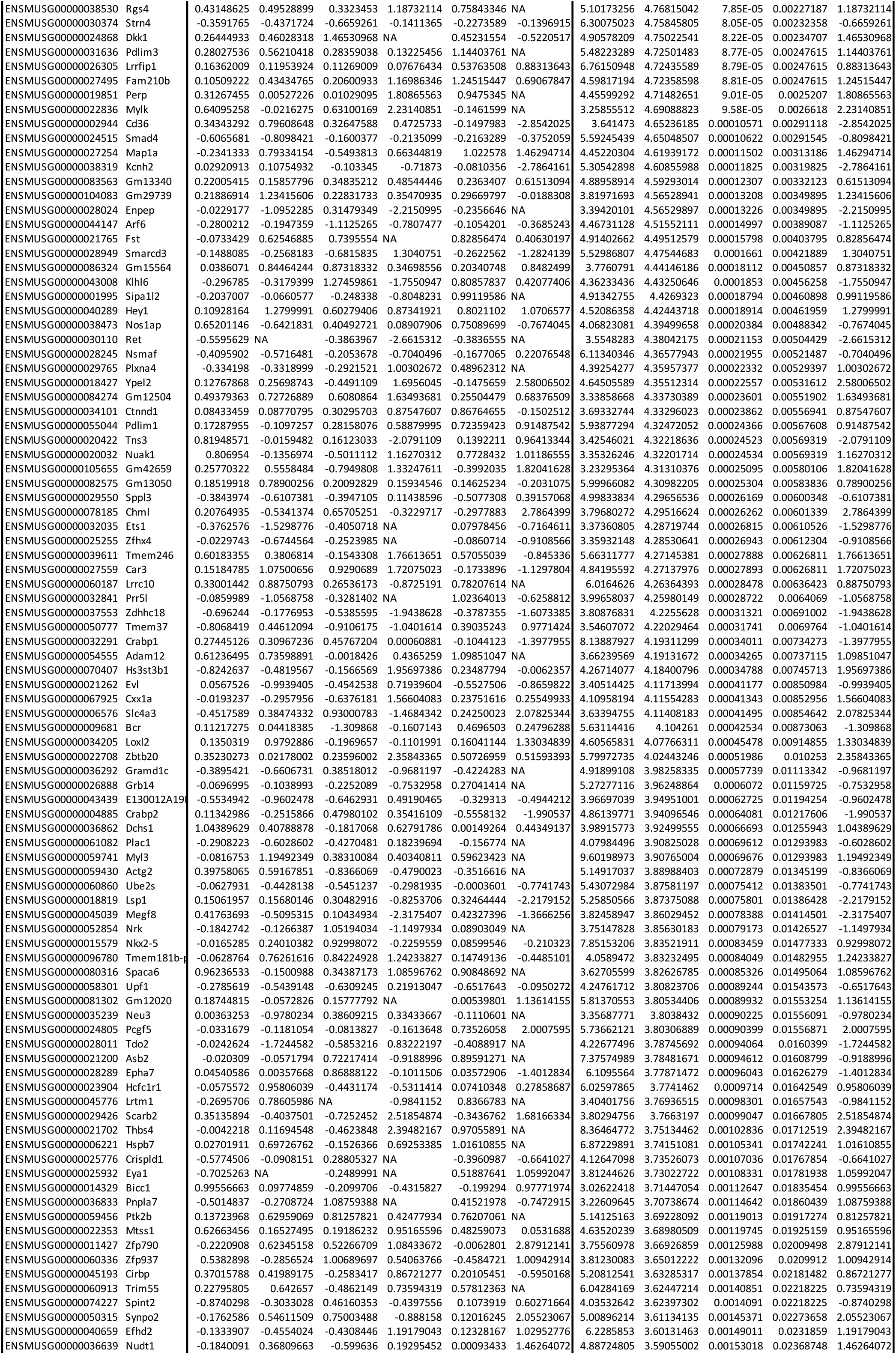
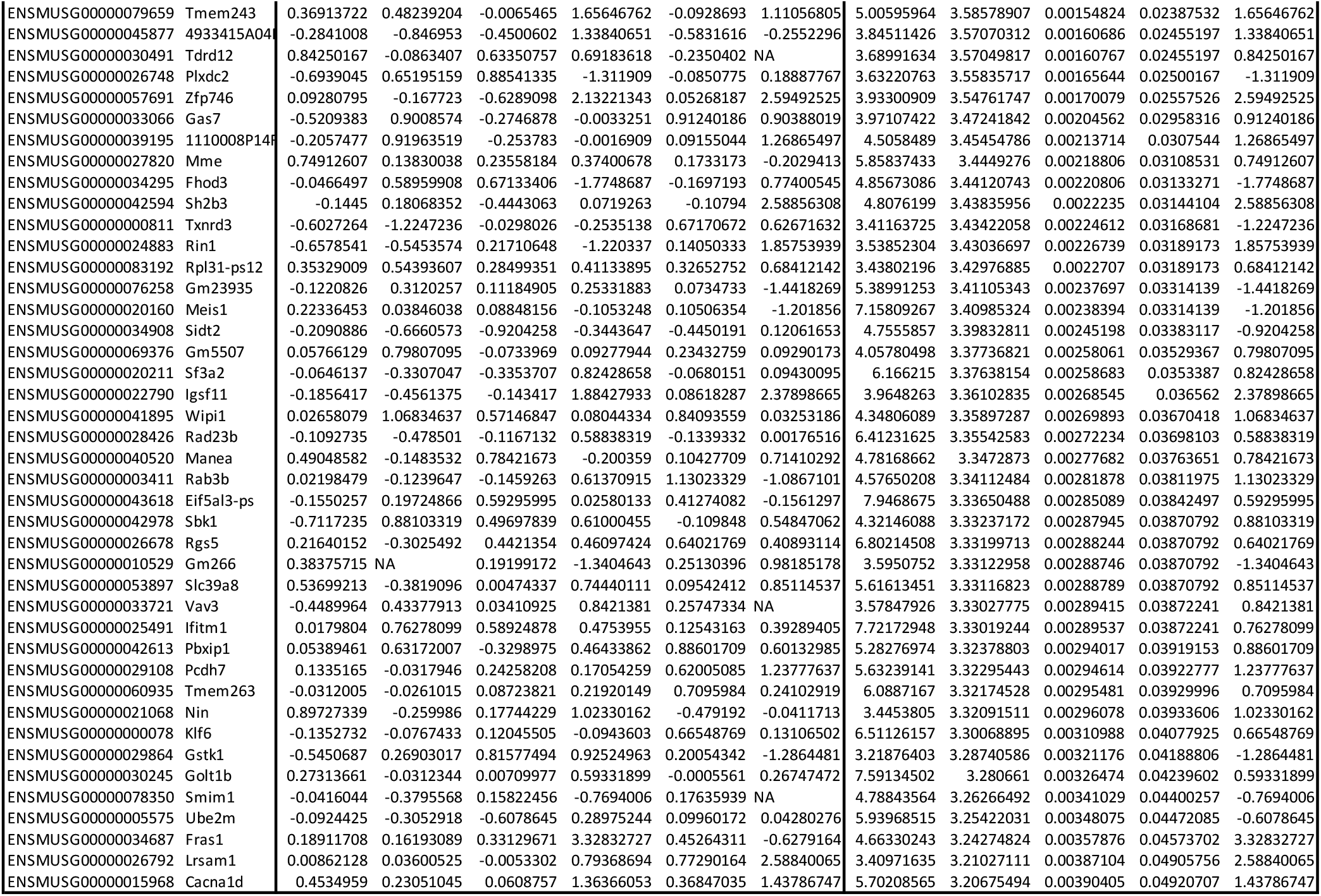
List of differentially expressed genes in left and right anterior mesoderm. Results reported for all genes considered significantly differentially expressed. The fold-change in each cluster is reported. Clusters with very low expression were filtered out and the fold-change is NA. A summary fold-change is reported (**logFC**) and corresponds to the largest (absolute) fold-change from all clusters. Genes are considered significantly DE if FDR < 0.05 **AND** they show an absolute fold-change for at least one cluster larger than 1.5.

## Notes

### Competing Interest Statement

The authors have declared no competing interest.

## REFERENCES

1. Little, R. B. & Norris, D. P. Right, left and cilia: How asymmetry is established. Semin Cell Dev Biol 110, 11– 18 (2021).

2. Collignon, J., Varlet, I. & Robertson, E. J. Relationship between asymmetric nodal expression and the direction of embryonic turning. Nature 1996 381:6578 381, 155– 158 (1996).

3. Lowe, L. A. et al. Conserved left–right asymmetry of nodal expression and alterations in murine situs inversus. Nature 1996 381:6578 381, 158–161 (1996).

4. Nonaka, S. et al. Randomization of left-right asymmetry due to loss of nodal cilia generating leftward flow of extraembryonic fluid in mice lacking KIF3B motor protein. Cell 95, 829–837 (1998).

5. Nonaka, S., Shiratori, H., Saijoh, Y. & Hamada, H. Determination of left–right patterning of the mouse embryo by artificial nodal flow. Nature 2002 418:6893 418, 96–99 (2002).

6. Meno, C. et al. Left–right asymmetric expression of the TGFβ-family member lefty in mouse embryos. Nature 1996 381:6578 381, 151–155 (1996).

7. Logan, M., Pagán-Westphal, S. M., Smith, D. M., Paganessi, L. & Tabin, C. J. The Transcription Factor Pitx2 Mediates Situs-Specific Morphogenesis in Response to Left-Right Asymmetric Signals. Cell 94, 307–317 (1998).

8. Ryan, A. K. et al. Pitx2 determines left–right asymmetry of internal organs in vertebrates. Nature 1998 394:6693 394, 545–551 (1998).

9. Yoshioka, H. et al. Pitx2, a bicoid-type homeobox gene, is involved in a lefty-signaling pathway in determination of left-right asymmetry. Cell 94, 299–305 (1998).

10. Piedra, M. E., Icardo, J. M., Albajar, M., Rodriguez-Rey, J. C. & Ros, M. A. Pitx2 Participates in the Late Phase of the Pathway Controlling Left-Right Asymmetry. Cell 94, 319–324 (1998).

11. Amand, T. R. S. et al. Cloning and expression pattern of chicken Pitx2: A new component in the SHH signaling pathway controlling embryonic heart looping. Biochem Biophys Res Commun 247, 100–105 (1998).

12. Kitamura, K. et al. Mouse Pitx2 deficiency leads to anomalies of the ventral body wall, heart, extra- and periocular mesoderm and right pulmonary isomerism. Development 126, 5749–5758 (1999).

13. Schweickert, A., Campione, M., Steinbeisser, H. & Blum, M. Pitx2 isoforms: involvement of Pitx2c but not Pitx2a or Pitx2b in vertebrate left–right asymmetry. Mech Dev 90, 41–51 (2000).

14. Franco, D. & Campione, M. The Role of Pitx2 during Cardiac Development: Linking Left–Right Signaling and Congenital Heart Diseases. Trends Cardiovasc Med 13, 157–163 (2003).

15. Tyser, R. C. v et al. Calcium handling precedes cardiac differentiation to initiate the first heartbeat. Elife 5, (2016).

16. Djenoune, L., Berg, K., Brueckner, M. & Yuan, S. A change of heart: new roles for cilia in cardiac development and disease. Nature Reviews Cardiology 2021 19:4 19, 211–227 (2021).

17. Desgrange, A., le Garrec, J. F., Bernheim, S., Bønnelykke, T. H. & Meilhac, S. M. Transient Nodal Signaling in Left Precursors Coordinates Opposed Asymmetries Shaping the Heart Loop. Dev Cell 55, 413-431.e6 (2020).

18. Esteban, I. et al. Pseudodynamic analysis of heart tube formation in the mouse reveals strong regional variability and early left–right asymmetry. Nature Cardiovascular Research 2022 1:5 1, 504–517 (2022).

19. le Garrec, J. F. et al. A predictive model of asymmetric morphogenesis from 3D reconstructions of mouse heart looping dynamics. Elife 6, (2017).

20. Behrens, A. N. et al. Nkx2-5 Mediates Differential Cardiac Differentiation Through Interaction with Hoxa10. Stem Cells Dev 22, 2211 (2013).

21. Tyser, R. C. V. et al. Characterization of a common progenitor pool of the epicardium and myocardium. Science 371, (2021).

22. Schweickert, A. et al. Left-asymmetric expression of Galanin in the linear heart tube of the mouse embryo is independent of the nodal co-receptor gene cryptic. Dev Dyn 237, 3557–3564 (2008).

23. Lawson, K. A. & Wilson, V. A Revised Staging of Mouse Development Before Organogenesis. Kaufman’s Atlas of Mouse Development Supplement Preprint at https://doi.org/10.1016/b978-0-12-800043-4.00003-8 (2016).

24. Amy Cooke, A. et al. The RNA-Binding Protein YBX3 Controls Amino Acid Levels by Regulating SLC mRNA Abundance SLC7A5 SLC3A2 Cell Reports The RNA-Binding Protein YBX3 Controls Amino Acid Levels by Regulating SLC mRNA Abundance. Cell Rep 27, 3097– 3106 (2019).

25. McGrath, J., Somlo, S., Makova, S., Tian, X. & Brueckner, M. Two populations of node monocilia initiate left-right asymmetry in the mouse. Cell 114, 61– 73 (2003).

26. Hummel, K. P. & Chapman, D. B. Visceral inversion and associated anomalies in the mouse. Journal of Heredity 50, 9–13 (1959).

27. Layton, W. M. Random determination of a developmental process: Reversal of normal visceral asymmetry in the mouse. Journal of Heredity 67, 336– 338 (1976).

28. Supp, D. M., Witte, D. P., Steven Potter, S. & Brueckner, M. Mutation of an axonemal dynein affects left–right asymmetry in inversus viscerum mice. Nature 1997 389:6654 389, 963–966 (1997).

29. Campione, M. et al. The homeobox gene Pitx2: mediator of asymmetric left-right signaling in vertebrate heart and gut looping. Development 126, 1225–1234 (1999).

30. Hoyt, P. R. et al. The Evi1 proto-oncogene is required at midgestation for neural, heart, and paraxial mesenchyme development. Mech Dev 65, 55–70 (1997).

31. Marques, S. et al. The activity of the Nodal antagonist Cerl-2 in the mouse node is required for correct L/R body axis. Genes Dev 18, 2342–2347 (2004).

32. Nakamura, T. et al. Fluid flow and interlinked feedback loops establish left–right asymmetric decay of Cerl2 mRNA. Nature Communications 2012 3:1 3, 1–13 (2012).

33. Sefton, M., Sánchez, S. & Nieto, M. A. Conserved and divergent roles for members of the Snail family of transcription factors in the chick and mouse embryo. Development 125, 3111–3121 (1998).

34. Murray, S. A. & Gridley, T. Snail family genes are required for left-right asymmetry determination, but not neural crest formation, in mice. Proc Natl Acad Sci U S A 103, 10300–10304 (2006).

35. Rago, L. et al. MicroRNAs Establish the Right-Handed Dominance of the Heart Laterality Pathway in Vertebrates. Dev Cell 51, 446-459.e5 (2019).

36. Balda, M. S., Garrett, M. D. & Matter, K. The ZO-1– associated Y-box factor ZONAB regulates epithelial cell proliferation and cell density. Journal of Cell Biology 160, 423–432 (2003).

37. Galli, D. et al. Atrial myocardium derives from the posterior region of the second heart field, which acquires left-right identity as Pitx2c is expressed. Development 135, 1157–1167 (2008).

38. Tyser, R. C. V. et al. Calcium handling precedes cardiac differentiation to initiate the first heartbeat. Elife 5, (2016).

39. Lawson, K. A. & Wilson, V. A Revised Staging of Mouse Development Before Organogenesis. in Kaufman’s Atlas of Mouse Development Supplement (2016). doi:10.1016/b978-0-12-800043-4.00003-8.

40. Picelli, S. et al. Full-length RNA-seq from single cells using Smart-seq2. Nat Protoc (2014) doi:10.1038/nprot.2014.006.

41. Choi, H. M. T. et al. Third-generation in situ hybridization chain reaction: Multiplexed, quantitative, sensitive, versatile, robust. Development (Cambridge) (2018) doi:10.1242/dev.165753.

